# Evaluating working memory representations during the attentional blink: A comparative modeling approach

**DOI:** 10.1101/2022.09.15.508098

**Authors:** Shuyao Wang, Aytaç Karabay, Elkan G. Akyürek

## Abstract

The nature of working memory (WM) limitations has been a topic of long-standing debate, with several models proposed to elucidate this issue. In this study, we conducted a systematic comparison of seven visual WM models to assess their ability to account for target consolidation during the attentional blink (AB). The AB phenomenon refers to where participants often fail to encode the second of two targets when there is a short time interval of ∼500 msec or less between them, providing an opportunity to evaluate commensurate WM limitations. Despite the growing consensus on the applicability of some WM models, such as the standard mixture model and the variable precision model, to the AB domain, no study has systematically evaluated these models in this context. We compared the performance of seven widely adopted visual WM models in four different AB datasets, drawn from three separate laboratories. We fitted each model and computed the Akaike Information Criterion (AIC) values at an individual level, across different conditions and experiments, based on which we compared the models. Slot-family models most often minimized AIC for second-target reports at short lags, while variable-precision models improved at longer lags and with color targets, indicating predominantly discrete consolidation during the AB, with feature- and lag-dependent graded components. These patterns imply that failure-to-encode (guessing) dominates over low-precision encoding, except when feature content or lag affords partial consolidation, refining theories of episodic tokenization and WM consolidation during the AB.

## 1. Introduction

Visual working memory (WM) is essential for information processing, yet it is challenging to encode all information completely. A substantial amount of research has focused on investigating the limitations of WM, which include storage limitations – the capacity of how many items can be encoded (Cowan, 2001; Miller, 1956; Pashler, 1988); but also processing limitations – the difficulties faced when processing and consolidating multiple items that appear either concurrently or in a rapid succession (Jolicœur & Dell’Acqua, 1998; Ricker & Cowan, 2014).

To investigate the limits of WM, an analysis of the kind of representation errors that occur in WM tasks has proven fruitful. Several WM models on the nature of these errors have been proposed. One perspective suggests that WM is comprised of a fixed, limited number of discrete “slots” dedicated to item storage, with large memory errors (i.e., those far away from the target value) occurring when items are not stored in WM, reflecting pure guessing (Alvarez & Cavanagh, 2004; Bays et al., 2009; Zhang & Luck, 2008). Another set of models proposes that WM is a continuous resource that can be distributed freely over items (Fougnie et al., 2012; van den Berg et al., 2012). According to this view, large memory errors arise when items are encoded with extremely low precision as opposed to high precision. Still other models focus on perceptual confusability and on the degree to which items are processed as an integrated ensemble. (Brady & Alvarez, 2011; Schurgin et al., 2020).

Although such model-based analyses of representation errors originate from, and have mainly been applied in, experimental tasks that target WM specifically, maintaining stimulus representations is critical to a myriad of cognitive functions, including selective attention, where target items have to be identified, selected, and reported on, sometimes also after a small delay. Through the application of model-based analyses, here we asked how targets are (mis-)reported in a temporal attention task, to assess the nature of representation during the attentional blink (AB) phenomenon.

The AB refers to the failure to identify the second (T2) of two briefly presented target stimuli, when the time interval (or lag) between them is within 200 – 500 msec (Broadbent & Broadbent, 1987; Raymond et al., 1992). Recent models of the AB suggest that the blink occurs due to the ongoing consolidation of the initial stimulus (Wyble et al., 2009, 2011; but see Zhao et al., 2024). During the consolidation of the first target (T1) into WM, attention is suppressed, which impairs the ability to identify new stimuli (e.g., T2). Event-related potential (ERP) studies provided evidence that the consolidation of T2 information is impaired, as shown by a decrease in the P3 component’s amplitude (Bourassa et al., 2015; Dell’Acqua et al., 2003; Vogel et al., 1998; Sergent et al., 2005; for a review see Zivony & Lamy, 2022). Further evidence for the involvement of WM in the emergence of the AB phenomenon has come from experiments showing that AB magnitude is increased when WM load is higher (Akyürek et al., 2007), which is also accompanied by reduced P3 amplitude (Akyürek et al., 2010).

Traditionally, AB research has utilized categorical items such as letters, numbers, or images in the rapid serial visual presentation (RSVP) paradigm (for a review, see Dux & Marois, 2009; Martens & Wyble, 2010). While this typical paradigm has proven to be an effective method for studying the AB itself, it presents inherent challenges in evaluating the quality of target representations maintained in WM during the AB window. In response to this challenge, recent studies have incorporated the continuous report task into the RSVP paradigm, as is also frequently utilized in the visual WM field (Bays et al., 2009; Fougnie & Alvarez, 2011; Kool et al., 2014; Oberauer et al., 2017; Zhang & Luck, 2008). During the task, participants are required to memorize targeted visual items, which are identified by a basic visual property that varies on a continuous, usually circular, dimension, such as color or orientation. In the reproduction stage, participants are then asked to recall the target on a continuous response scale, such as recreating its orientation by rotating a Gabor (Wilken & Ma, 2004). Consequently, the performance of a continuous report task is not measured in a binary format, but rather as the divergence between the target value and the corresponding response, making it possible to quantify the target representations stored in WM. Thus, by combining the AB paradigm with continuous report tasks, we can measure the structure of T2 errors, and assess whether target report failures arise from the rate of all-or-none losses or from (overall) degradation in precision.

Several previous studies have explored this approach. For instance, Asplund et al. (2014) applied a mixture-model analysis (Zhang & Luck, 2008) to data obtained by explicitly asking participants to recall their perception of T2 along a continuous circular dimension. Through this approach, the authors were able to quantify the precision of T2 representations and the proportion of guess responses for T2. Their findings indicated that AB affects T2 perception in an all-or-none manner, with an increased percentage of random guess trials at shorter lags, while lag had no effect on T2 precision.

In a similar vein, Karabay et al. (2022) employed hierarchical Bayesian estimation to explore the variability in the precision and guess rate parameters of the standard mixture model at different time intervals between two targets. Their study showed that T2 awareness during the AB can be both gradual and discrete. Specifically, when the identification of the first target (T1) required the focus of attention on a single spatial location, T2 awareness was discrete, with the guess rate varying significantly; on the contrary, when T1 identification induced a spatial spread of attention, T2 awareness was gradual, with variations in precision. Sy et al. (2021) utilized both the standard mixture and variable precision models, and also found that the temporal loss of T2 information during the AB might be either gradual or discrete. Specifically, when two targets (T1 and T2) shared a visual feature, their results indicated a gradual loss of T2 precision, with the variable precision model providing a better fit to the data. In contrast, they observed a discrete loss of T2 information when participants had to switch their attention to different target features (i.e., color for T1 and orientation for T2), and in this case the mixture model performed better than the variable precision model in terms of fitting the data. While the primary objective of Sy et al. was to discern whether perception during the attentional blink is discrete or graded, it is worth noting that their results present mixed results in terms of model performance.

Despite these previous efforts to apply WM models to AB tasks, a comprehensive account of target representation during the AB has not yet emerged. In large part, this may be due to the variable outcomes, which could be due to differences in task paradigms and sample variability that have not been assessed systematically. Moreover, most comparisons among WM models have primarily focused on the standard mixture and variable precision models, potentially overlooking other relevant components which could affect WM representations. For example, the influence of non-target items proposed by the ensemble integration model (Brady & Alvarez, 2011) may be relevant in the context of the AB. This hypothesis is supported by Akyürek and Hommel (2005; Akyürek et al., 2012), who discovered that temporal integration occurs with successive targets in RSVP. This suggests that representational errors during the blink may also reflect interactions with temporally adjacent items in the stream.

Against these open questions, the current study examines how WM representations are shaped under temporal constraints, by applying a broader set of WM models to continuous report data gathered from multiple AB tasks. We start by selecting seven commonly used visual WM models: the standard mixture model, the slot model, the slots plus resource model, the swap model, the ensemble integration model, and two versions of the variable precision model. These models fundamentally diverge in how they interpret memory errors in the continuous reproduction task. We also considered the target confusability competition model (Schurgin et al., 2020), due to its new perspective on modeling working memory representations, but it was ultimately excluded from our analysis. The target confusability competition model quantifies psychophysical similarity within a given feature space (e.g., color), elucidating how familiarity propagates within that specific stimulus field. However, we found that it produced unstable fits across features under RSVP timing depending on the specific feature, likely due to psychophysical similarity functions inadequately estimated under RSVP. To ensure the generalizability and robustness of our results, we analyzed continuous report data from three previously conducted AB studies by distinct research groups (Asplund et al., 2014; Karabay et al., 2022; Tang et al., 2020), together with data from a new experiment we have undertaken ourselves.

## 2. Method

### 2.1. Model Details

#### Slot model family: Models with limited number of WM slots

The family of slot models, including the standard mixture, slot and slots plus resource models is built upon the core assumption that WM consists of a limited number of discrete slots for storing visual representations (Cowan, 2001; Luck & Vogel, 1997). If the target object has been stored in one of these slots, its information is maintained in WM. In contrast, if the item has not been stored in any of these slots, then no information about it remains in WM.

##### Standard mixture model (StM)

The StM model (Zhang & Luck, 2008) proposes a mixture distribution for responses. When items are successfully stored into WM, the corresponding response values tend to cluster around the actual value, thereby forming a Von Mises distribution. Conversely, when item information is not available in WM, the reported values are expected to be random, conforming to a uniform distribution. The probability of this guessing is denoted by the height of this uniform distribution. Taken together, the response error distribution in the StM model is given by:

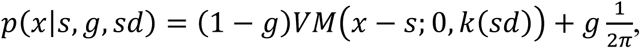

where *x* is the reported value, *s* is the target value. The model parameter *g*, or the guess rate, indicates the proportion of trials where subjects are presumed to make random guesses. The von Mises distribution is expressed as:

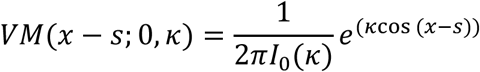

In this formula, *κ*, the concentration parameter of the Von Mises distribution, signifies the precision of memory representations. It is analogous to the standard deviation *sd* in the normal distribution. *I*_0_(*κ*) is the modified Bessel function of the first kind of order 0, which ensures that the total probability density integrates to 1.

##### Slot model

The Slot model (Cowan, 2001; Luck & Vogel, 1997), which serves as the classic representation of this family, posits that there is a fixed number of storage slots in WM. When the number of items to be remembered, known as the set size, exceeds the maximum number of available slots, any excess items are disregarded. Consequently, in the Slot model, the guess rate (*g*) is correlated with the maximum number of slots. The response error distribution is formally given by:

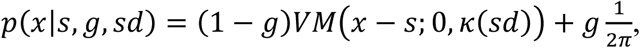

The guess rate *g* depends on the model parameter, *capacity* (*K*), according to the following relationship. When *K* ≥ *N* (set size), all items can be remembered, and the guess rate would equal zero. When *K* < *N*:

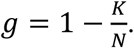

##### Slots plus resource model (SR)

The SR model (Awh et al., 2007; Zhang & Luck, 2008), like previous models, hypothesized a fixed, finite capacity of WM wherein a certain number of items can be stored. What distinguishes the SR model is its unique approach to WM resource allocation. Specifically, most WM resources are preferentially assigned to a probed item, with only a minimal amount for other items that are considered less relevant. The SR model proposes that when the number of items is fewer than the slots available, the surplus resources improve the precision of the stored items. In such instances, we anticipate significant variance in the precision (standard deviation, *sd*) of WM, depending on the set size. Formally, the response error distribution is given by:

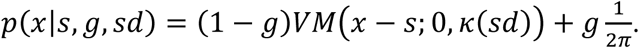

Here, the *sd* is extrapolated using the SR model’s parameters, *bestSD* and the number of probed items *n*, as follows:

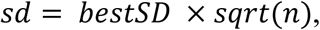

where *bestSD* denotes the precision when all WM resources are used for storing a single item, and *n* is the number of items that can be stored in WM, depending on the *capacity K* and set size *N*:

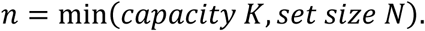

The guess rate is linked to the *capacity K* in the same way as in the Slot model.

It should be noted that the key difference between the StM model and the other two slot models (the Slot and SR models) is that the latter explicitly proposes hypotheses about the effect of set size on error distribution. In the AB paradigm, the set size is typically considered to be 2, which may result in similar performance across these models due to the consistent set size. However, we also consider the condition where T1 is incorrectly reported (see Section 2.4, Data Preparation, for a detailed explanation). We expect that this condition will reveal differences in the models’ performance as the set size changes.

#### The variable precision models: Models that assume WM consists of a resource pool

The variable precision models (Fougnie et al., 2012; van den Berg et al., 2012) propose that WM resources are allocated to items in a continuous and variable manner, which causes the precision of WM representation to vary across items and trials. Although the variable precision models and the SR model both suggest that WM is not (entirely) quantized, there are two primary distinctions between them. First, the variable precision models define WM limitations by the quality of the representation, as opposed to the number of items memorized. This is in contrast to the SR model, which puts emphasis on the item count. Second, the variable precision models imply a variability of WM resources across trials and items. The SR model, in contrast, implies an equal level of precision across all items and trials, when the set size is fixed.

##### Variable precision model (VP)

In the VP model proposed by van den Berg et al. (2012), item information does not undergo a total loss. Instead, the quality of the item representation may severely degrade, resulting in extremely low precision. Therefore, the response error distribution is given by:

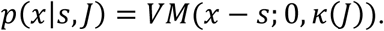

Here, each item is encoded with precision *J*, which is defined as the Fisher information. The concentration parameter *κ* is related to *J* through the following relationship:

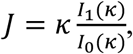

where *I*_0_ represents the modified Bessel function of order zero and *I*_1_ represents the modified Bessel function of order one.

The core hypothesis in the VP model is that the precision *J* varies independently across items and trials. Furthermore, the precision *J* itself follows a specific distribution as a random variable. Van den Berg et al. (2012) further assume this specific distribution of precision *J* to be Gamma with mean precision *J* and scale parameter *τ*:

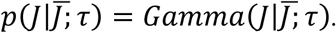

Therefore, the response error distribution of the VP model is a mixture of an infinite set of Von Mises distributions:

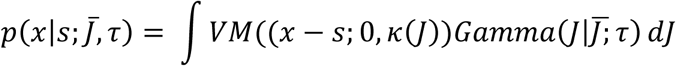

##### Variable precision model with guess rate (VPG)

In the current study, we also employed another version of the VP model with an extra *guess rate* parameter (Fougnie et al., 2012). Mathematically, the response distribution is then given by:

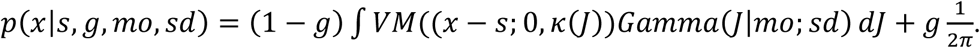

Thus, the VPG model has three free parameters: guess rate (*g*), the mode (*mo*) and the standard deviation (*sd*) of the Gamma distribution of precision *J*.

#### Interference models: Models that assume there are interaction between items

##### Ensemble integration model (EnsInt)

The aforementioned models suggest that each item is stored in WM independently, which implies there is no interaction between these items. However, Brady and Alvarez (2011) demonstrated that the representation in WM is not only affected by the item itself, but also by the integration of information of all the stimuli that are displayed. Their results showed that the reported size of a target item was shifted toward the mean size of the items with the same color, as well as the mean size of all items displayed. The EnsInt model implemented in the current study is a simplified version of the Brady & Alvarez (2011) model, which introduces a parameter that represents the bias towards the mean value of all distractors. Since the first target is the only other relevant item in typical continuous-report AB tasks, we used T1’s value for this parameter, slightly simplifying the model. Formally, the error distribution of the EnsInt model is given by:

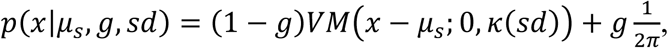

where *μ*_*s*_ refers to the biased mean value, which is estimated based on the differences between the target (T2) and the other item (T1).

##### Swap model

This model can be considered as an extension of the StM model. In addition to the two components of the response error distribution, namely a uniform distribution that indicates the proportion of random-guessing trials and a Von Mises distribution that represents the memory precision, Bays et al. (2009) introduced a third source of error: The probability of incorrectly reporting non-target items. Consequently, the Swap model has a three-component structure, including precision, guess rate, and swap rate (estimated by the measurement of the distance between reported values and non-target items):

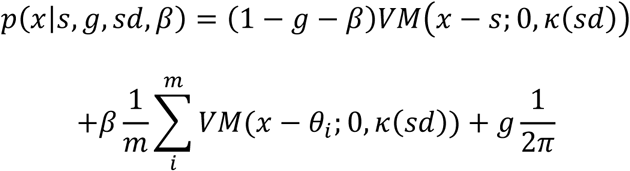

where *β*, the swap rate, denotes the probability of incorrectly reporting the non-target items (distractors, i.e., T1) and where {*θ*_1_, *θ*_2_, … *θ*_*m*_} are the values of a set of *m* distractors.

### 2.2. Datasets

As summarized in Table 1, we obtained three existing datasets from three separate laboratories (Asplund et al., 2014; Karabay et al., 2022; Tang et al., 2020), and added a fourth, new dataset in the present paper (Wang et al.).

**Table 1.** Details of the data sets.

| Datasets | Exp | SOA (ms) | Lag | T1 feature | T2 feature | subject number |
| --- | --- | --- | --- | --- | --- | --- |
| *Asplund et al., 2014* | Exp 1 | NA - | 1, 2, 4, 8 | Categorical | Color (1°-360°) | 25<sup>1</sup> |
| *Tang et al., 2020* | Exp 1 | 50, 240,<br>360, 600,<br>720 | 1, 2, 3, 5,<br>7 | Orient. (1°-180°) | Orient. (1°-180°) | 22 |
|  | Exp 2 | 360, 720 | 3, 7 | Orient. (1°-180°) | Orient. (1°-180°) | 23 |
| *Karabay et al., 2022* | Exp 1A | 250, 800 | NA | Orient. (1°-180°) | Orient. (1°-180°) | 23 |
|  | Exp 1B | 250, 800 | NA | Orient. (1°-180°) | Orient. (1°-180°) | 21 |
|  | Exp 2A | 250, 800 | NA | Color (1°-360°) | Color (1°-360°) | 26 |
|  | Exp 2B | 250, 800 | NA | Color (1°-360°) | Color (1°-360°) | 27 |
|  | Exp 3 | 300, 800 | 3, 8 | Orient. (1°-180°) | Orient. (1°-180°) | 24 |
|  | Exp 4 | 300, 800 | 3, 8 | Orient. (1°-180°) | Orient. (1°-180°) | 24 |
|  | Exp 5 | 300, 800 | 3, 8 | Orient. (1°-180°) | Orient. (1°-180°) | 23 |
| *Wang et al. data* | Exp 1 | 90, 300, 800 | 1, 3, 8 | Orient. (1°-180°)<br>or Color (1°-180°) | Orient. (1°-180°)<br>or Color (1°-180°) | 28 |

*Note.* Exp = Experiment; Orient. = Orientation; Wang et al. data refers to the new dataset collected in the present study.

**Asplund et al. (2014).** The first dataset was obtained from Experiment 1 in Asplund et al. (2014), where square-shaped targets were embedded in circle-shaped colored distractors. T1 was filled with either black or white, while the T2 color was randomly chosen from 180 equiluminant colors. T2 appeared at the first, second, fourth, or eighth position (labeled as Lag 1, Lag 2, Lag 4, and Lag 8) following T1. The stimulus duration for each participant was set by a staircase method and resulted in an average value of 150 ms (*SD* = 10 ms). At the end of the stimulus stream, participants were asked to reproduce T2 using a color wheel, and to report whether the first target was black or white.

**Tang et al. (2020).** The second dataset was retrieved from two behavioral experiments in the AB study by Tang et al. (2020), where a RSVP task was again employed. Specifically, there were 20 Gabor patches, and the orientation of the Gabors were integer angles ranging from 0 to 179 degrees, without replacement. Participants were instructed to memorize two targets (high-frequency Gabors) among distractors (low-frequency Gabors) and reproduce them at the end of each trial. Each target or distractor appeared for 40 ms, and the next item followed after a blank interval of 80 ms. The time interval between two targets was manipulated by presenting different numbers of distractors in-between (lag). In Experiment 1, there were five lag conditions (1, 2, 3, 5, 7), while in Experiment 2, there were two lag conditions (3, 7).

**Karabay et al. (2022).** Karabay et al. (2022) used two different paradigms, dwell time (DT) and RSVP, both with continuous report tasks, to examine the nature of awareness in the AB. We acquired seven datasets from their experiments. Experiment 1A, 1B, 2A, and 2B were run with the DT paradigm. Both targets were orientation gratings (in Experiment 1A and 1B) or colors (in Experiment 2A and 2B). T1 and a distractor were presented, at the same time, above or below the fixation dot, at an equal distance. The display of T2 was the same as T1, except the location changed to the left or right side of the fixation dot. On each trial, T1 and T2 target arrays were followed by a mask. The stimulus-onset asynchrony (SOA) between T1 and T2 was set to 250 ms and 800 ms. The RSVP paradigm was used in Experiment 3, 4, and 5, in which the targets were always orientation gratings. The number of distractor items between two targets was 2 (Lag 3, 300 ms SOA) or 7 (Lag 8, 800 ms SOA). In Experiment 3, the two targets appeared among a stream of distractors, similar to the design in Tang et al. (2020). In Experiment 4, there were simultaneously two streams of stimuli at the left and right sides of the fixation dot. In Experiment 5, T1 was presented within a single stream of distractors, but split into a dual stream with the presentation of T2, as in Experiment 4.

### 2.3. Wang et al. data

In the new experiment, we manipulated feature-dimension changes between targets in the AB task (same or different), hypothesizing that matching target feature-dimensions might cause mutual bias in target reports, which might affect the model selection. Like the other studies, this experiment also featured continuous report of targets.

#### 2.3.1. Participants

The minimum sample size of 24 subjects was determined by means of a G-power analysis (Faul et al., 2007), with the following parameters: significance level (*α* = .05), beta-level (*β* = .2), and a medium to large effect size (Cohen’s *dz* = .6). Twenty-eight undergraduate students (seventeen females and eleven males) from the University of Groningen were recruited in exchange for course credits (mean age = 19.6, range = 18-27). All participants reported normal or corrected-to-normal visual acuity and no color blindness. Informed consent forms and instructions were given to the participants prior to participation. Prior to its execution, ethical approval was obtained from the ethical committee of the Psychology Department of the University of Groningen (Approval code PSY-1819-S-0209). The study was conducted in accordance with the Declaration of Helsinki (2008).

#### 2.3.2. Apparatus and stimuli

Participants were individually seated on a desk chair in sound-attenuated lab chambers, at a viewing distance of approximately 60cm from a 22” CRT monitor (Iiyama MA203DT). The screen was set at a 1280 x 1024-pixel resolution, 16-bit color depth, and a 100Hz refresh rate. The trials were prepared and run in OpenSesame 3.2 (Mathôt et al., 2012) with the PsychoPy back-end (Peirce, 2007; 2009), under the Microsoft Windows 7 operating system. All the stimuli were displayed in the center of the screen on a grey background (RGB: 128,128,128), as shown in Figure 1A. There were two kinds of target stimuli in the RSVP stream (Figure 1B). The first kind of target consisted of orientation gratings with a spatial frequency of 1.8 cycles/degree of visual angle, presented within a circle of 2.2° of visual angle. The orientation of the gratings was chosen randomly from the range of 0°-179°. The other kind of target consisted of circles (2.2° of visual angle) filled with one of 360 colors, which were chosen randomly from the HSL color spectrum. The distractors were grey-scale mosaic images (2.2° × 2.2° of visual angle), retrieved from Karabay et al. (2022).

**Figure 1.**
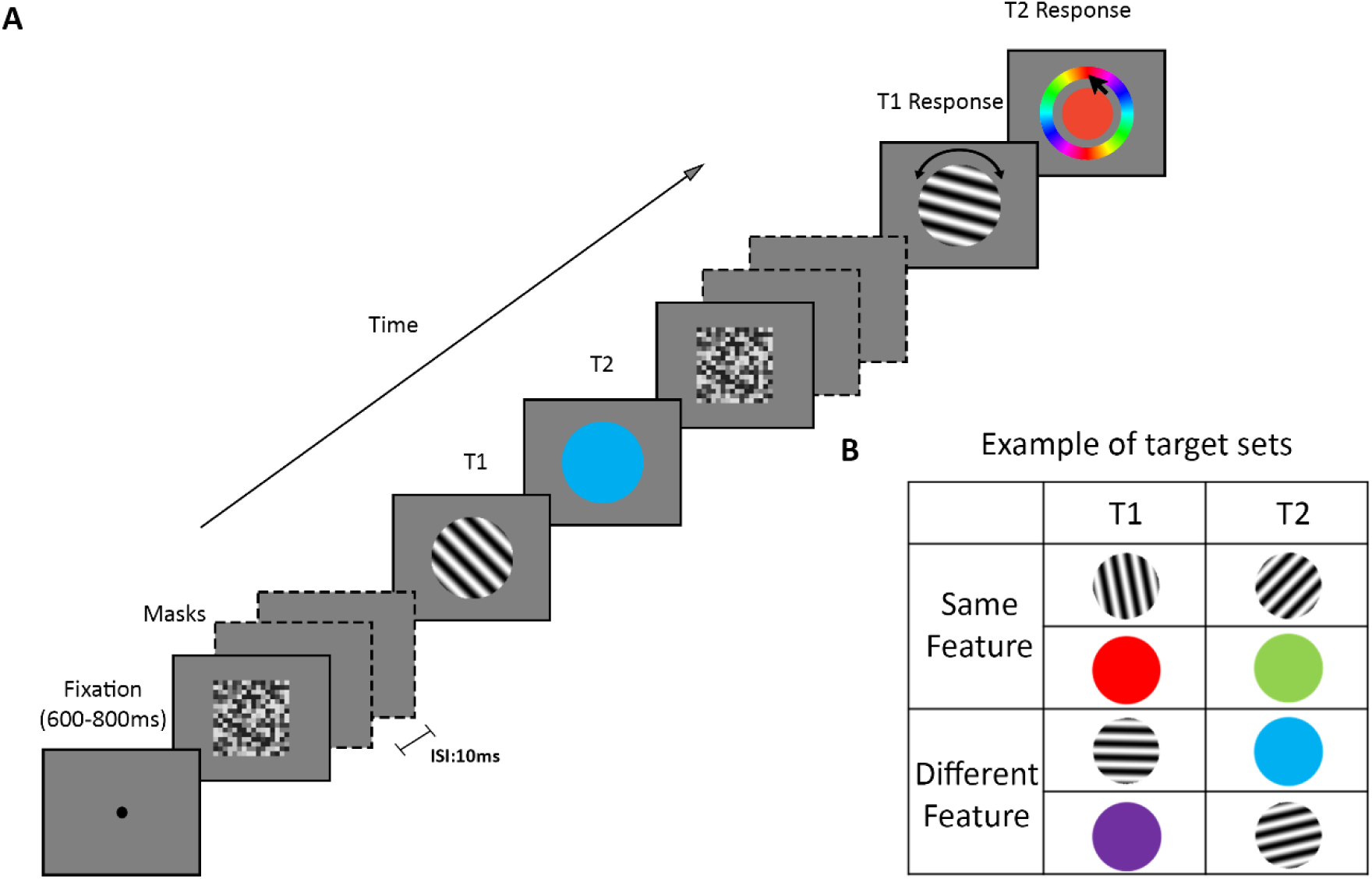
Illustration of the design of the newly conducted experiment. *Note.* (**A**) An example trial of the dual-target RSVP task at Lag 3. In this instance, the two targets have different feature-dimensions. (**B**) Examples of possible targets in all combinations and orders of the two targets.

#### 2.3.3. Procedure

A 2 (Target feature-dimension: same or different) x 3 (Lag: 1, 3, 8) factorial design was implemented in this experiment. Two practice blocks, each including 16 trials, were administered at the beginning, which were excluded from the analysis. After that, there were 540 experimental trials evenly distributed into 15 blocks. Each trial started with a fixation dot that randomly lasted for 600-800ms. There were 18 items within the RSVP stream. Each item was shown for 80 ms, separated by a 10 ms inter-stimulus interval (ISI). T1 was randomly presented at the fifth, sixth, or seventh position of the stream. T2 was presented consecutively (Lag 1), after two items (Lag 3), or after seven items (Lag 8), following T1. T1 and T2 could have either the same feature-dimension (color or orientation), or different feature-dimensions. At the end of the stream, after a blank interval of 500ms, two response prompts were displayed successively.

Participants were asked to reproduce the two targets in the correct order. They were asked to reproduce the target orientation by rotating a probe grating, or to reproduce the target color by choosing the exact color from a color wheel. Responses were collected with a standard mouse. Trial-based feedback was provided only in practice trials: A happy smiley was shown for correct trials, and an unhappy smiley for incorrect trials. Responses close to target values (below 22.5° reproduction error) were considered correct responses for feedback. In the experimental blocks, the average performance on T1 and T2 were displayed at the end of each block. During the experiment, participants could have a self-paced break between blocks. The whole session took about 90 minutes to complete.

### 2.4. Data Preparation

As indicated, all the data sets we gathered for analysis were obtained from continuous-report AB tasks. We reorganized the datasets to form a uniform structure, as per the following procedure:

1. For each trial, response error for each target (T1 and T2) was calculated by subtracting the response value from the actual one.
2. To avoid cases of trivial near-identity and ambiguous swap/integration, trials in which the absolute deviations between T1 and T2 values were less than 22.5 degrees were discarded.
3. Response errors were limited to a range from −90° to 90° for orientations and from - 180 to 180 for colors.
4. Performance of T2 was evaluated in two distinct conditions:

a. T1 correct Condition (T2|T1) – This condition is often employed in AB studies. In this condition, we included only the trials in which the absolute error of T1 was less than or equal to 22.5 degrees for orientations (45 degrees for colors).
b. T1 incorrect condition – This condition included the trials in which the absolute error of T1 exceeded 22.5 degrees for orientations (45 degrees for colors). This condition served as a proxy for reduced effective load, but since it is not fully equivalent to a true single-item set size, our conclusions using this proxy are bracketed accordingly.

Given that the set size remained constant across all our data sets, we took into account that some models (e.g., the SR model) incorporate set size for parameter estimation. Hence, we also analyzed T2 performance when T1 was reported incorrectly, as this represents a scenario where the set size equals one. Although this condition does not strictly correspond to a true single set size condition, it does provide a valuable approximation of model performance when the memory load is set to one, while the typical T2|T1 performance analysis evaluates the models’ performance when the memory load is two.

### 2.5. Model fitting

We employed maximum likelihood estimation (MLE) as our chosen approach to fit all of the aforementioned models to our data sets. The general logic of MLE is to maximize the likelihood function to find the specific parameter value, with which the chosen probabilistic model is most “likely” to generate the observed data. Specifically, for each subject and condition within a single experiment, we carried out model fitting using the built-in MLE function in MemToolbox (Suchow et al., 2013). This allows us to obtain the best estimates of the model parameters at the individual level (See Supplementary Materials A for details of parameter estimates).

### 2.6. Model comparison

In terms of model comparison and selection, it is important to strike a balance between model flexibility and overfitting. Although flexible models can accommodate a wide range of datasets by introducing additional parameters, they may lack supporting evidence and can lead to overfitting (Ziegel, 2003). To assess the fitness of the models and determine the best model for describing our data, we used the Akaike Information Criterion (AIC; Akaike, 1974) as a common penalized model comparison measure.

The AIC values were calculated based on the natural logarithm of the likelihood function, also known as the log-likelihood. Specifically, the AIC was calculated using the formula:

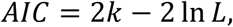

where *k* represents the number of parameters, and *L* denotes the maximized value of the likelihood function. At the individual level, we obtained the AIC values using the model comparison function implemented in MemToolbox (Suchow et al., 2013; memtoolbox.org). Lower AIC values indicate a better fit of the model to the data. By subtracting the lowest AIC value (corresponding to the most likely model for each subject) from the AIC values of other models, we obtained the relative AIC values at the subject level.

Additionally, to extend our analysis to the group level, we investigated the model performance against group data. In detail, we first averaged the relative AIC values across all subjects for each experimental condition (e.g., lag). Second, we ranked the models according to their AIC values for each subject and presented the average rankings across all subjects, separately for each condition. This allowed us to gain insights into the overall model performance and make meaningful comparisons among the models within each condition.

### 2.7. Simulation analysis

Although the AIC indicates how well a model fits the experimental data compared to other models, it does not provide a comprehensive assessment of each model’s absolute performance. To gain a more detailed understanding of how well each model captures different aspects of the data, we conducted a simulation analysis. This involved generating simulated data sets from these assessed models and comparing the model predictions to the actual T2 errors in the observed data.

To perform the simulations, we utilized the most probable parameter values from the model fitting procedure described earlier (See *2.5. Model fitting*). These parameters were applied to their respective models for each subject and each condition. We then generated simulated datasets with the same number of data points as those collected from each subject in the corresponding condition. Subsequently, we calculated the summary statistics of T2 errors separately for the simulated data, and the actual data gathered from different studies. This simulation procedure was implemented with a custom-made function in MATLAB, based on the *SampleFromModel* function in Memtoolbox.

## 3. Results

### 3.1. Asplund et al. (2014)

#### 3.1.1. Model comparison

Figures 2 and 3 present the model comparison analysis for the Asplund et al. (2014) data. Since Asplund et al.’s data consisted of entirely different T1 and T2 stimulus, the EnsInt model and the Swap model were excluded as they both need a third parameter for the similarity of two targets that was absent here.

**Figure 2.**
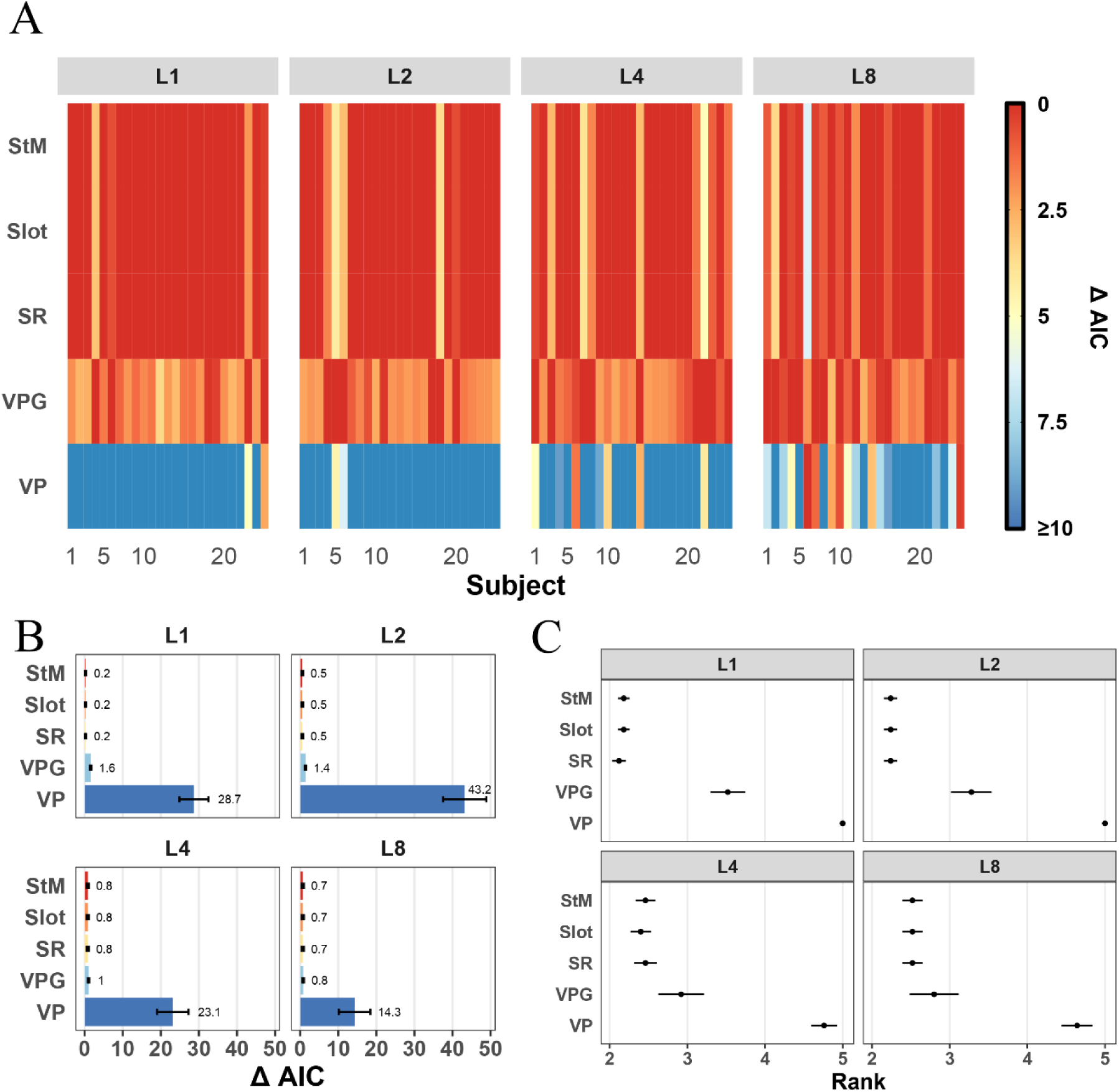
Model comparison for T2 performance when T1 was correct (T2|T1). *Note.* (**A**) Model comparison results at the subject level, using the data from Asplund et al. (2014). Each column represents a subject, and each row represents a specific model. Each panel represents a lag condition. L1 = Lag 1; L2 = Lag 2; L4 = Lag 4; L8 = Lag 8. Cell color indicates each model’s relative AIC value. Larger relative AIC values mean worse performance. (**B**) Relative AIC values compared to that of the best-performing model, averaged across all subjects. Error bars indicate standard error. (**C**) Model ranking. The ranks were obtained by averaging the per-subject ranks for each specific lag. Larger ranks mean worse performance. The model abbreviations are mentioned in the Method (see *2.1. Model details*).

**Figure 3.**
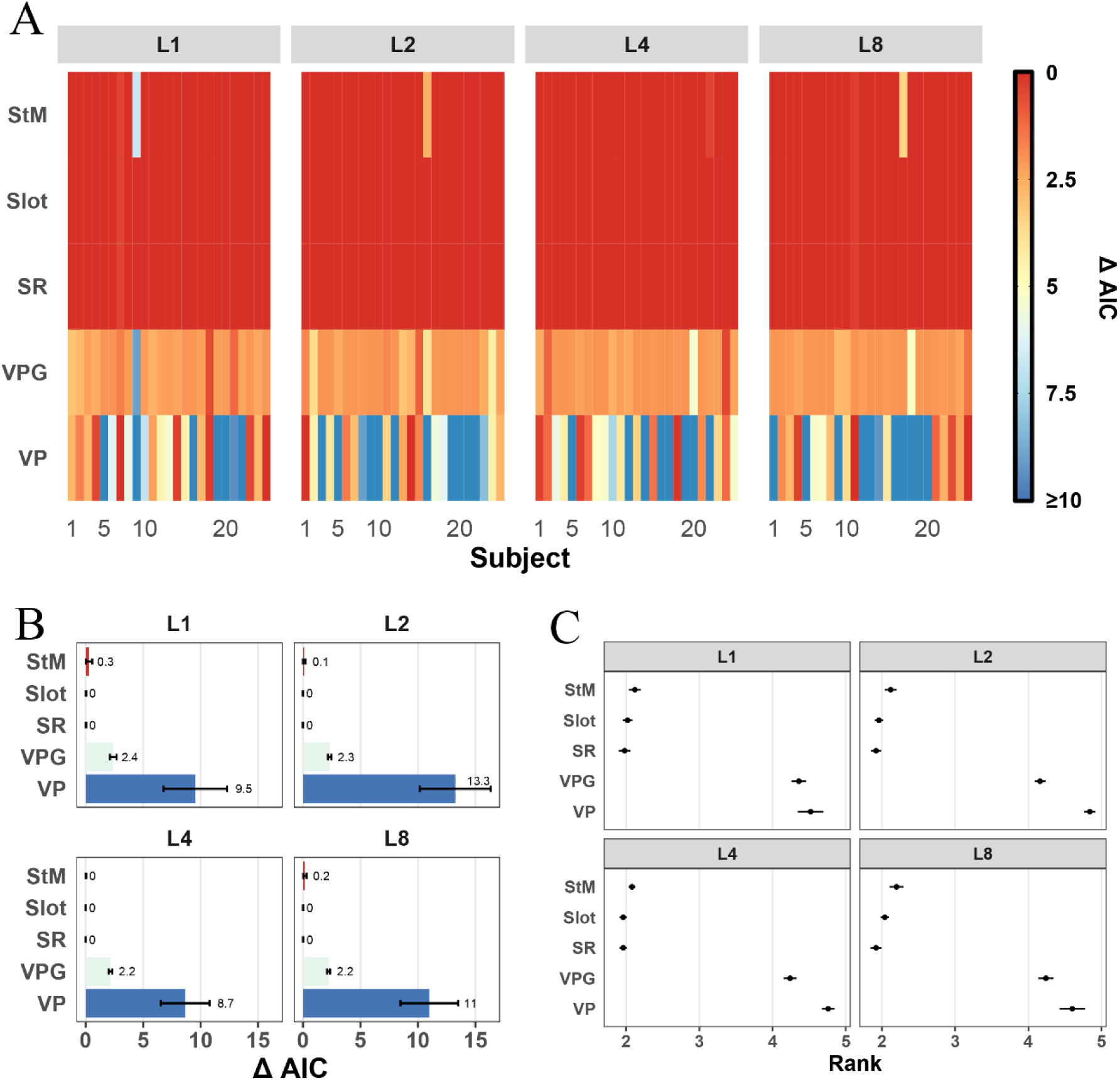
Model comparison for T2 performance when T1 was incorrect. *Note.* (**A**) Model comparison results at the subject level, using the data from Asplund et al. (2014). (**B**) Relative AIC values, averaged across all subjects for each lag. (**C**) Model ranking at the group level. Figure conventions follow Figure 2.

Figure 2 shows the results of the T1 correct condition, that is, T2|T1 performance. At the individual level (Figure 2A), the StM, Slot, and SR models consistently exhibited the best fit across most subjects’ data. In terms of the VP model, it consistently showed the poorest performance across almost all subjects, particularly at Lag 1 and 2^2^. Interestingly, the VPG model fitted better than the VP model. As the lag increased, the performance of the StM, Slot, and SR models declined for some subjects, while the performance of the VPG model improved.

To further assess model performance at the group level, we summarized the individual model fitness values in several ways. First, we calculated the AIC values relative to the best-performing model and then derived their average values across all subjects for each lag (as illustrated in Figure 2B). All models performed commendably across all lags, except for the VP model, which exhibited smaller relative AIC values as the lag increased. Next, we ranked the models for each subject based on their AIC values and averaged these ranks over all subjects for each lag (Figure 2C). This pattern closely mirrored the model performance depicted in Figure 2B. The StM, Slot, and SR models consistently achieved the lowest ranks (and thus, superior) across all lags. The VPG model, while ranking higher (thus poorer) at Lag 1 and 2, improved its performance at longer lags (Lag 4 and 8). This is consistent with the model comparison results of the VPG model at the individual level, with more subjects getting lower AICs when lag increased. Moreover, the consistency of the general model performance across different lags indicated that the AB itself (in particular at Lag 2) did not change the shape of the response error distributions, compared to other lags.

Figure 3 presents the results of model comparison for T2 performance in the T1 incorrect condition. The model comparison pattern bears a resemblance to that in the T1 correct condition, observable at both individual and group levels. Having superior performance across all lags were the StM, Slot, and SR models. In contrast, the VP model consistently demonstrated poor performance.

#### 3.1.2. Simulation analysis

We compared experimentally observed and simulated T2 accuracy for each respective lag. Figure 4 displays the average T2 accuracy across all subjects in both T1 correct and incorrect conditions. Of particular interest was the response error of the second target when the time interval between the first and second target fell between 200 – 500 ms (primarily at Lag 2), to get an idea if these models can actually describe the data well within the AB window. In general, the pattern of the simulated data was more or less similar to that of the actual data, except for the simulated data obtained from the VP model. The VP model had difficulty in mimicking a ‘blink’ shape in the error, presenting relatively similar T2|T1 errors across lags and generally lower values compared to the actual data and other model simulations In terms of T2 performance when T1 was incorrect, there was still a large divergence between the simulated data of the VP model and the actual data.

**Figure 4.**
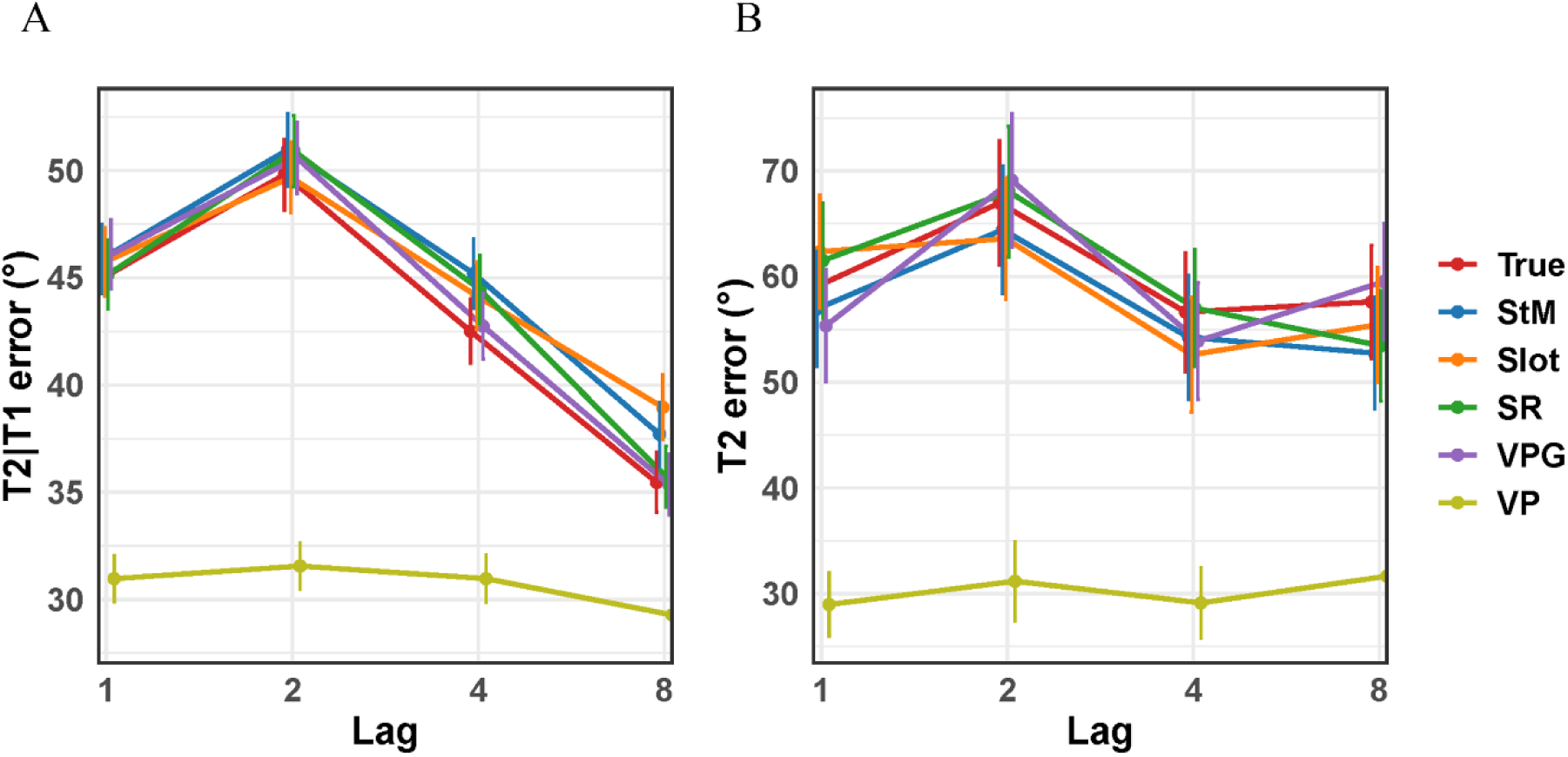
Comparison of model simulations and experimental data based on T2 accuracy. *Note.* Model simulation results using the data from Asplund et al. (2014). (**A**) Average T2 errors in the T1 correct condition. (**B**) Average T2 errors in the T1 incorrect condition. True = experimentally observed data.

### 3.2. Tang et al. (2020)

#### 3.2.1. Model comparison^3^

Figure 5 depicts the model comparison results for T2|T1 performance. At the individual level, the StM, Slot and SR models performed almost identical (Figure 5A). At Lag 1, 2 and 3, these three models showed a good fitness for some individuals, while they behaved poorly for the others (ΔAICs ≥ 16). As lag increased, model performance was improved, and the number of bad individual fits decreased. However, it can clearly be seen that the Swap model consistently showed excellent, if not best, fitness across all subjects and persistently at all lags. It is also worth noting that the number of participants for which the Swap model was the winning one dropped as a function of lag. At Lag 1, 2, and 3, the EnsInt model was the best model for a subset of subjects. The extent of preference for the EnsInt model decreased at Lag 5 and 7, where the EnsInt model showed a relatively consistent and moderate performance across all subjects. The VPG model’s performance was only better than its variant, the VP model, for some subjects at short lags (Lag 1, 2, and 3). However, at long lags (Lag 5 and 7), its performance was improved for most subjects. In contrast, the VP model was always the worst fitting one for all subjects at all lags, except for some individuals at Lag 5 and 7. In Experiment 2, there were two lag conditions (Lag 3 and Lag 7). The results presented a similar pattern to Experiment 1 with regard to lag. It was noteworthy that the Swap model was again the winning model for a subset of subjects at Lag 3, but only for two subjects at Lag 7. The swap model’s superiority at Lag 1–3 and decline at Lag 5–7 is consistent with the occurrence of temporal integration / order reversals at short lags, and reduced nontarget-influence at longer lags.

**Figure 5.**
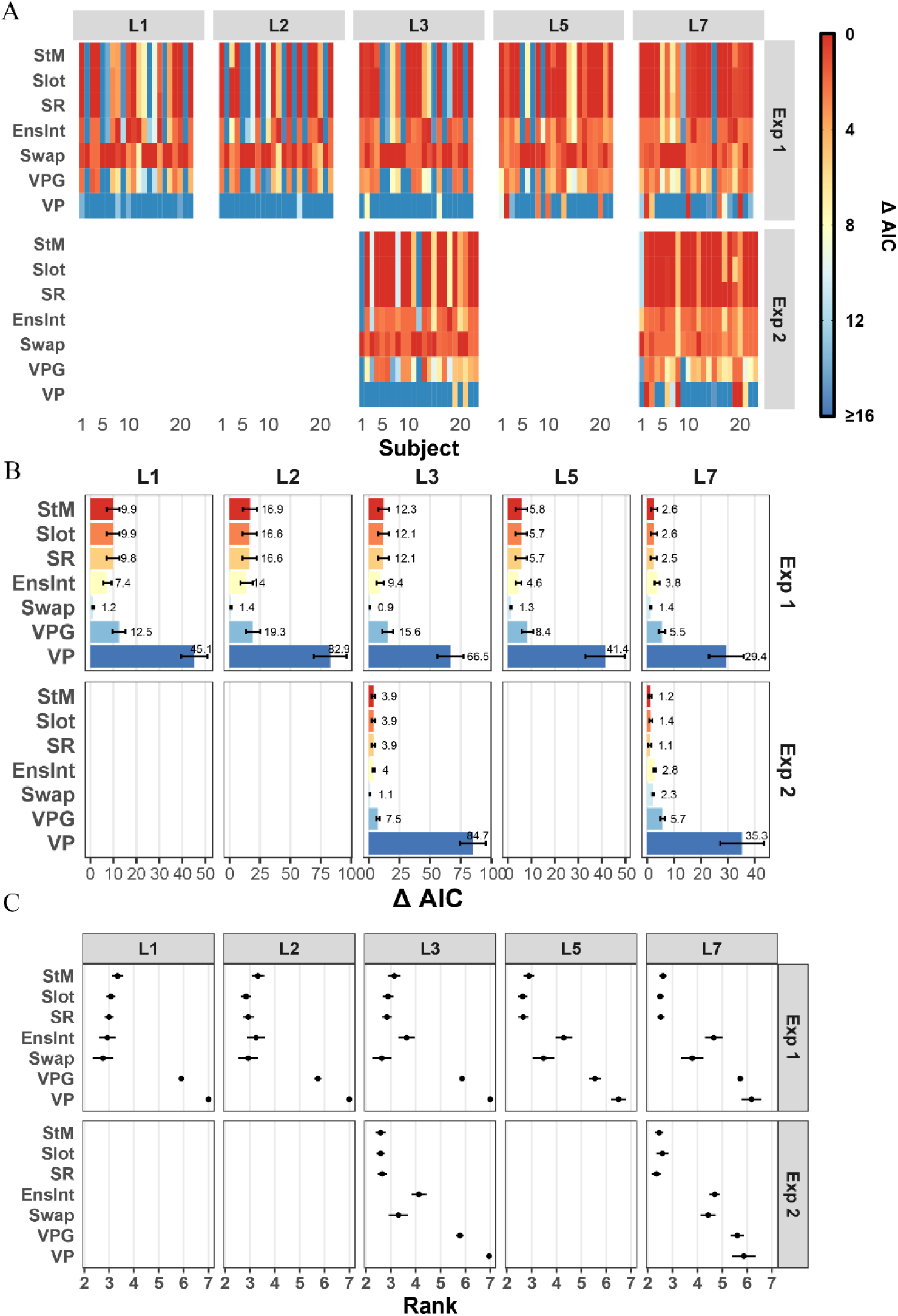
Model comparison for T2 performance when T1 was correct (T2|T1). *Note.* (**A**) Model comparison at the subject level, using the data from Tang et al. (2020). L1 = Lag 1; L2 = Lag 2; L3 = Lag 3; L5 = Lag 5; L7 = Lag7. Exp = Experiment. (**B**) Relative AIC values compared to that of the best-performing model, averaged across all subjects. (**C**) Average model ranking. Figure conventions follow Figure 2.

**Figure 6.**
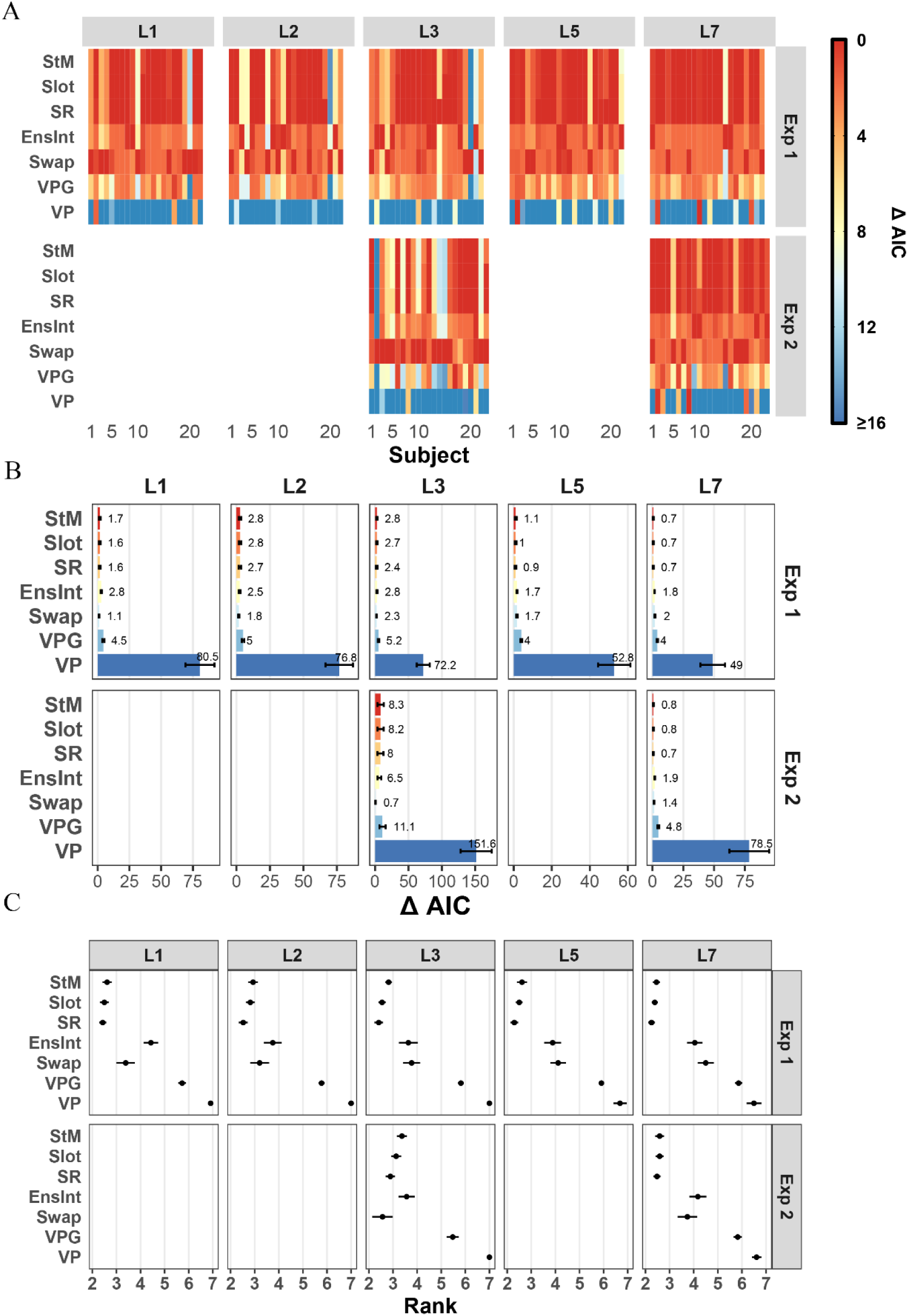
Model comparison for T2 performance in the T1 incorrect condition. *Note.* (**A**) Model comparison results at the individual level, using the data from Tang et al. (2020). (**B**) Relative AIC values, averaged across all subjects for each lag and each experiment, respectively. (**C**) Average model ranking. Figure conventions follow Figures 2 and 5.

The averaged relative AIC values (as displayed in Figure 5B) indicated a lag-dependent change in the models’ fitness. During Experiment 1, the StM, Slot, SR and EnsInt models reached their peak relative AIC values at Lag 2, followed by a decrease at Lag 5 and 7. The VP and VPG models exhibited a similar pattern. Remarkably, the Swap model consistently registered the smallest ΔAIC values at all lags, maintaining a value of less than 2 points. This finding is coherent with existing AB theories such as the eSTST (Bowman & Wyble, 2007; Wyble et al., 2009), and with the notion of temporal integration (Akyürek & Hommel, 2005). These suggest that targets are bound to same episodic event if they are successively shown, possibly when the second target is processed (tokenized) more quickly than the first one. Then, timestamps of the targets might be confused, which causes T2 to be reported as T1 and vice versa. For Experiment 2, the Swap model recorded the lowest mean ΔAIC at Lag 3 (1.1), but a slightly higher value (2.3) at Lag 7, at which the StM, Slot and SR models took over the leading positions. Generally, the comparison results from Experiment 2 displayed a similar pattern to that observed in Experiment 1.

The average model ranking at the group level is shown in Figure 5C. For Experiment 1, at Lag 1, 2 and 3, the StM, Slot, SR, EnsInt and Swap models dominated the top ranks with minimal divergence between them. At Lag 5 and 7, however, the EnsInt and Swap models experienced an increase in ranks (indicating poorer performance) compared to the leading models (the StM, Slot and SR). The VP and VPG models, on the other hand, consistently ranked last. What stood out in this comparison was the differences in rank across different lags for the EnsInt and Swap models in Experiment 1, a feature not observed in the other models. In Experiment 2, the StM, Slot and SR models achieved the smallest ranks (thus, superior performance) at both Lag 3 and 7.

We observed that when T1 was incorrect, the performance of the models exhibited a more uniform pattern across subjects and lags in both Experiment 1 and Experiment 2. Evidently, the VP model underperformed at both the individual and group level, whereas other models presented either good or moderate fitness. Moreover, the divergences among the top-performing models were fairly minimal.

#### 3.2.2. Simulation analysis

Figure 7 displays the summary statistics for simulated T2 errors from seven models alongside the experimentally observed T2 errors from Tang et al.’s study (2021). It is evident that there was a considerable deviation between the simulated data from the VP model and the other models, compared to the clustering of the simulated data from these latter models around the experimental T2 errors. The differences between these models (except the VP model) and the experimental data were relatively narrow. However, it’s worth noting that in Experiment 1 and the T1 correct condition, the EnsInt model generated a distinct T2 response shape, with T2|T1 errors continually decreasing as lag increased (Figure 7A), while the True data set exhibited its highest T2|T1 errors at Lag2 (the AB effect). The discrepancy indicates a potential mismatch between the EnsInt model’s simulation and the actual experimental T2 responses.

**Figure 7.**
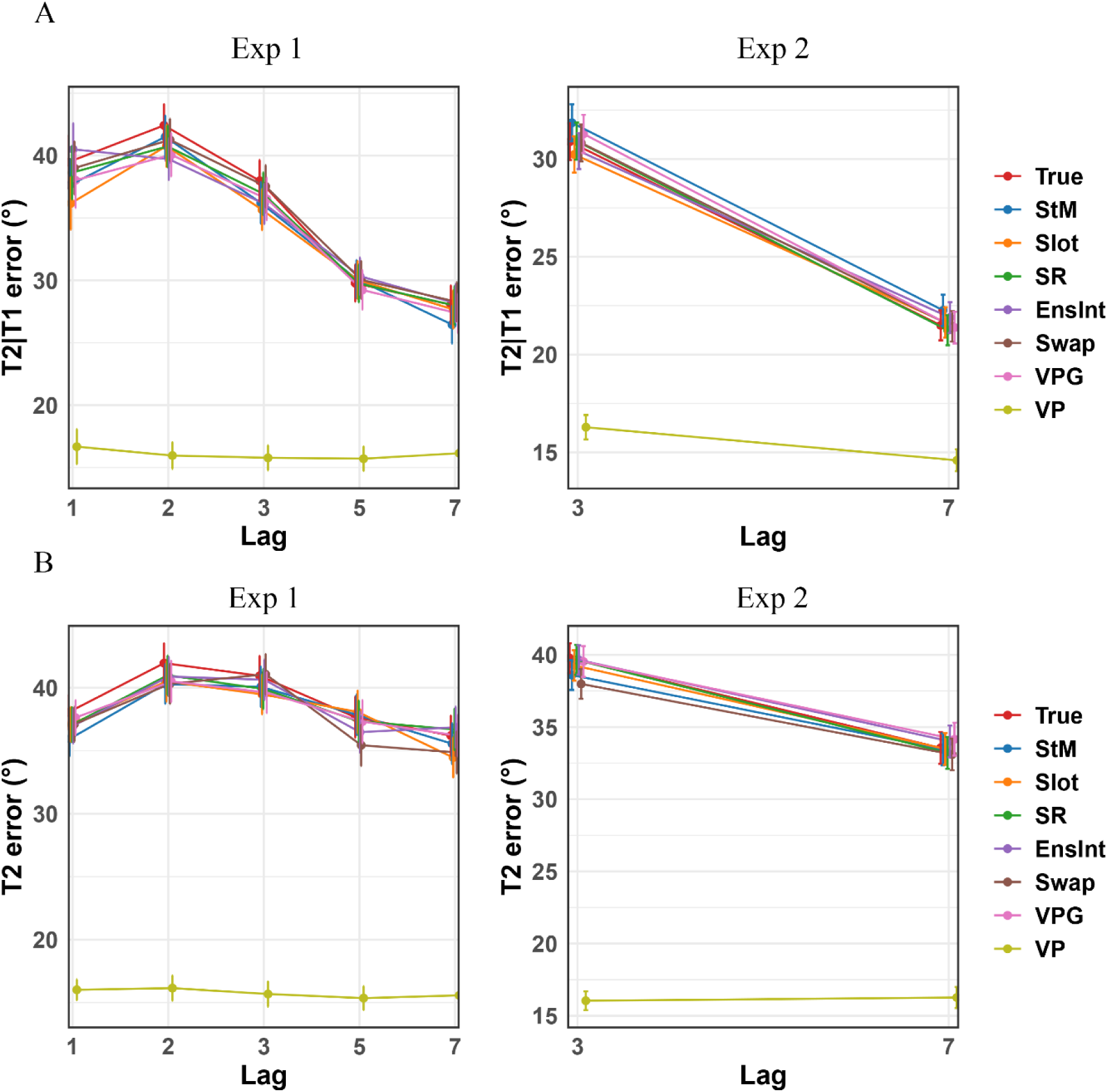
Comparison of model simulations and experimental data based on T2 accuracy. *Note*. Model simulation results using the data from Tang et al. (2020). Exp = Experiment. (**A**) Average T2 errors in the T1 correct condition. (**B**) Average T2 errors in the T1 incorrect condition. True = experimentally observed data. See Method for model abbreviations.

### 3.3. Karabay et al. (2022)

#### 3.3.1. Model comparison

Figure 8 presents the model comparison results for T2|T1 performance. When comparing these results across the various experiments within Karabay et al.’s (2022) study, the most apparent observation was that the models’ goodness-of-fit in Experiment 2 was clearly different from those in other experiments, which was evident at both the subject and group level.

**Figure 8.**
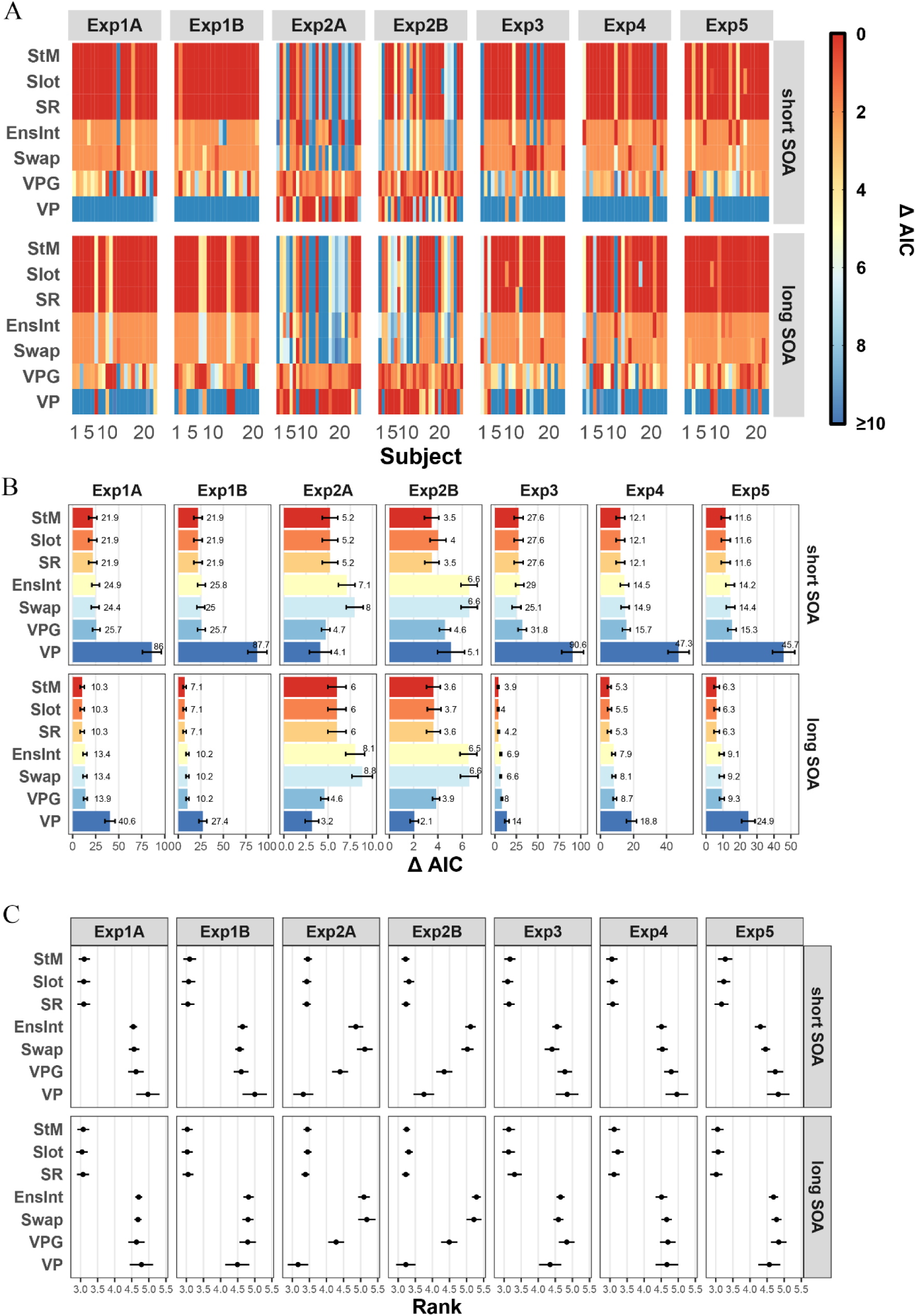
Model comparison for T2 performance when T1 was correct (T2|T1). *Note.* (**A**) Model comparison at the subject level, using the data from Karabay et al. (2022). Exp = Experiment; SOA = stimulus onset asynchrony. (**B**) Relative AIC values compared to that of the best-performing model, averaged across all subjects. (**C**) Average model ranking. Figure conventions follow Figure 2.

Delving into the results at the subject level, in Experiment 2A and 2B, where the targets had only a color feature, the VP and VPG models exhibited a good fit for most subjects at two distinct SOAs. Particularly at long SOA, the VP model outperformed all of its competitors for over half of the subjects. Conversely, the EnsInt and Swap models produced the least accurate data fit, maintaining either relatively higher or the highest ΔAIC values across all subjects. Intriguingly, here the StM, Slot, and SR models showed two extremes of fitness, either being the best or the worst.

In the results of the other experiments, a similar pattern of model fitness is observable. In detail, the StM, Slot, and SR models maintained the lowest ΔAIC values for the majority of subjects; the EnsInt and Swap models had relatively higher ΔAIC values. Additionally, although the VPG model’s fitness was the poorest for a subset of subjects, it demonstrated a moderate performance for the others, while the VP model scored the highest ΔAIC values for a majority of individuals.

In line with the results at the subject level, the VP and VPG models showed superior fitness at the group level in Experiment 2A and 2B, compared to other experiments, as depicted in Figure 8B. Furthermore, the range of mean ΔAIC values in Experiment 2A and 2B was markedly narrower than those observed in the other experiments. In terms of the model ranking (Figure 8C), the StM, Slot and SR models consistently claimed the leading positions in all experiments, with the exception of Experiment 2A and 2B. It is also worth noting that the EnsInt and Swap models ranked last in these two experiments.

Figure 9 presents the model comparison results for T2 performance when T1 was incorrect. Generally, the models demonstrated a relatively consistent performance across all subjects at the individual level. In addition to this, the pattern of model ranking, as inferred from their averaged ΔAIC values and their average ranks, mostly reflected those observed in the T1 correct condition. Yet, there were certain differences in comparison to the T1 correct condition. For example, although the VP and VPG models presented their best fit in Experiment 2A and 2B compared to other experiments, they still occupied the bottom ranks in Experiment 2A and 2B.

**Figure 9.**
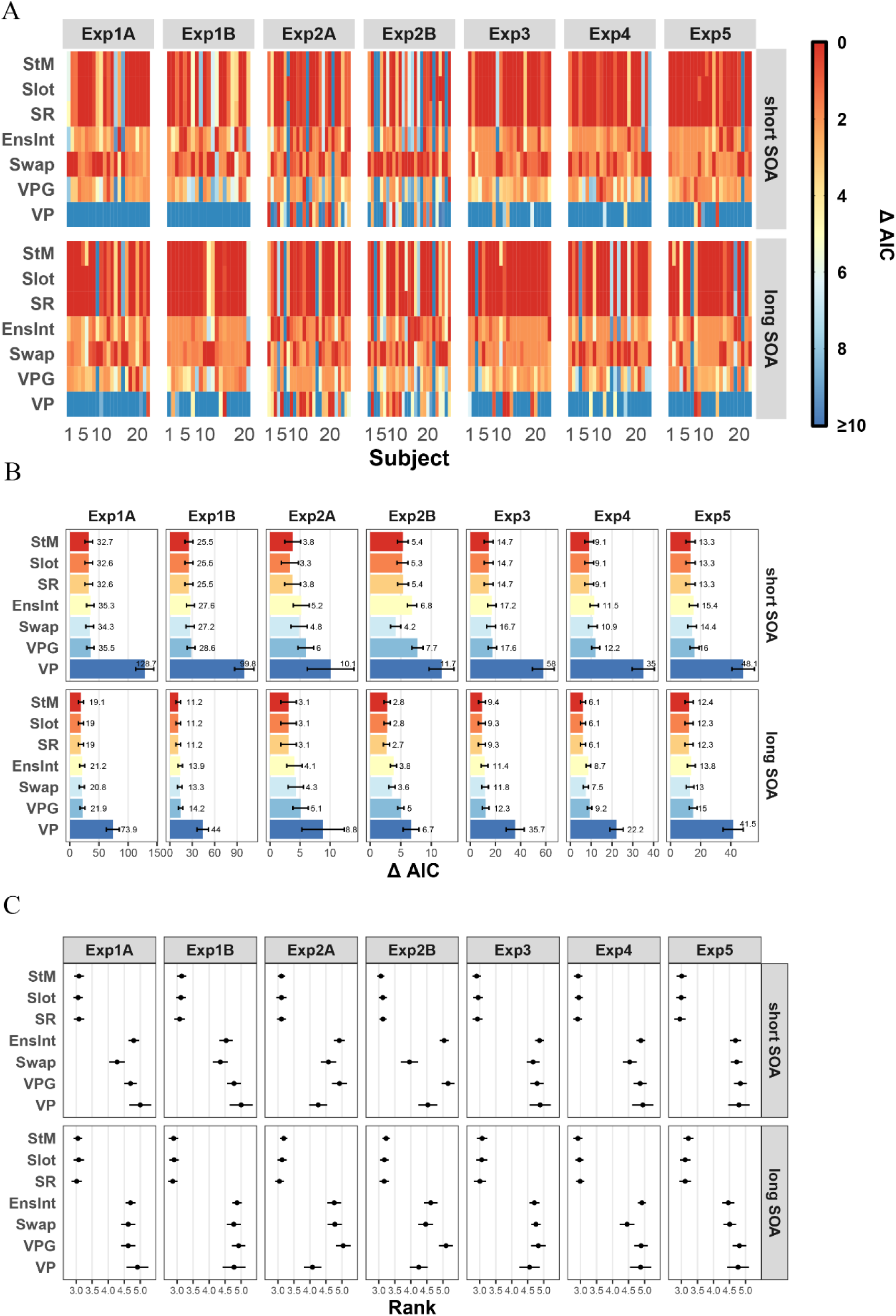
Model comparison for T2 performance when T1 was incorrect. *Note.* (**A**) Model comparison at the subject level, using the data from Karabay et al. (2022). Exp = Experiment; SOA = stimulus onset asynchrony. (**B**) Relative AIC values compared to that of the best-performing model, averaged across all subjects. (**C**) Average model ranking. Figure conventions follow Figure 2.

#### 3.3.2. Model Simulation

Figure 10 shows a comparison between the experimental data and the simulated data derived from all seven models. A notable discrepancy was again observed between the VP model and the other seven models. Specifically, most of the models (except for the VP model) provided simulations that were closely aligned with the experimental T2 response errors, regardless of whether T1 was correct or incorrect. Interestingly, in the T1 correct condition (Figure 10A), when the targets were colors (in Exp2A and 2B), the VP model’s pattern closely mirrored that of the other models, presenting a similar trend of T2 error across two distinct SOAs (or Lags). However, this good performance of the VP model was not observed in the T1 incorrect condition (Figure 10B).

**Figure 10.**
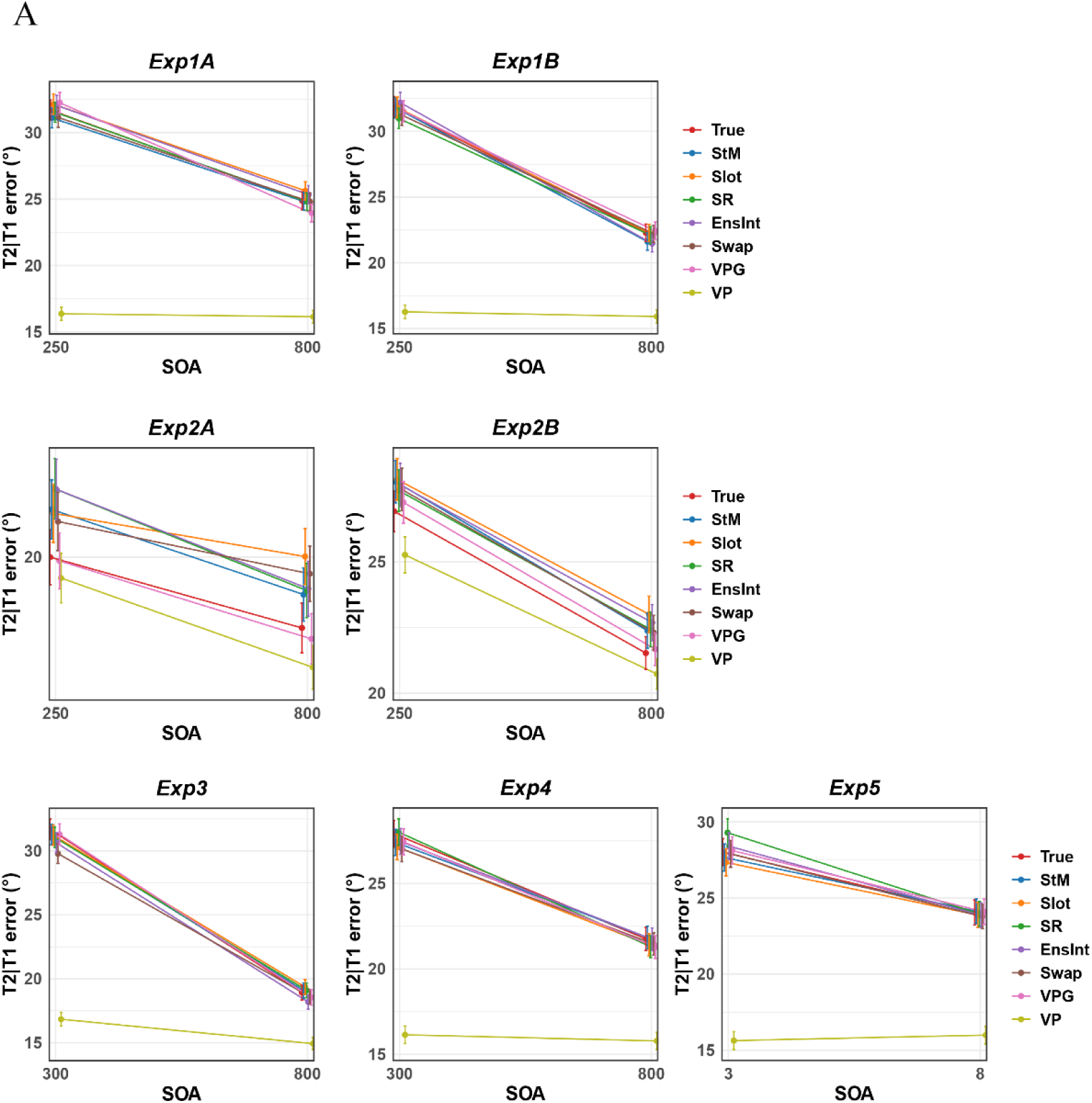
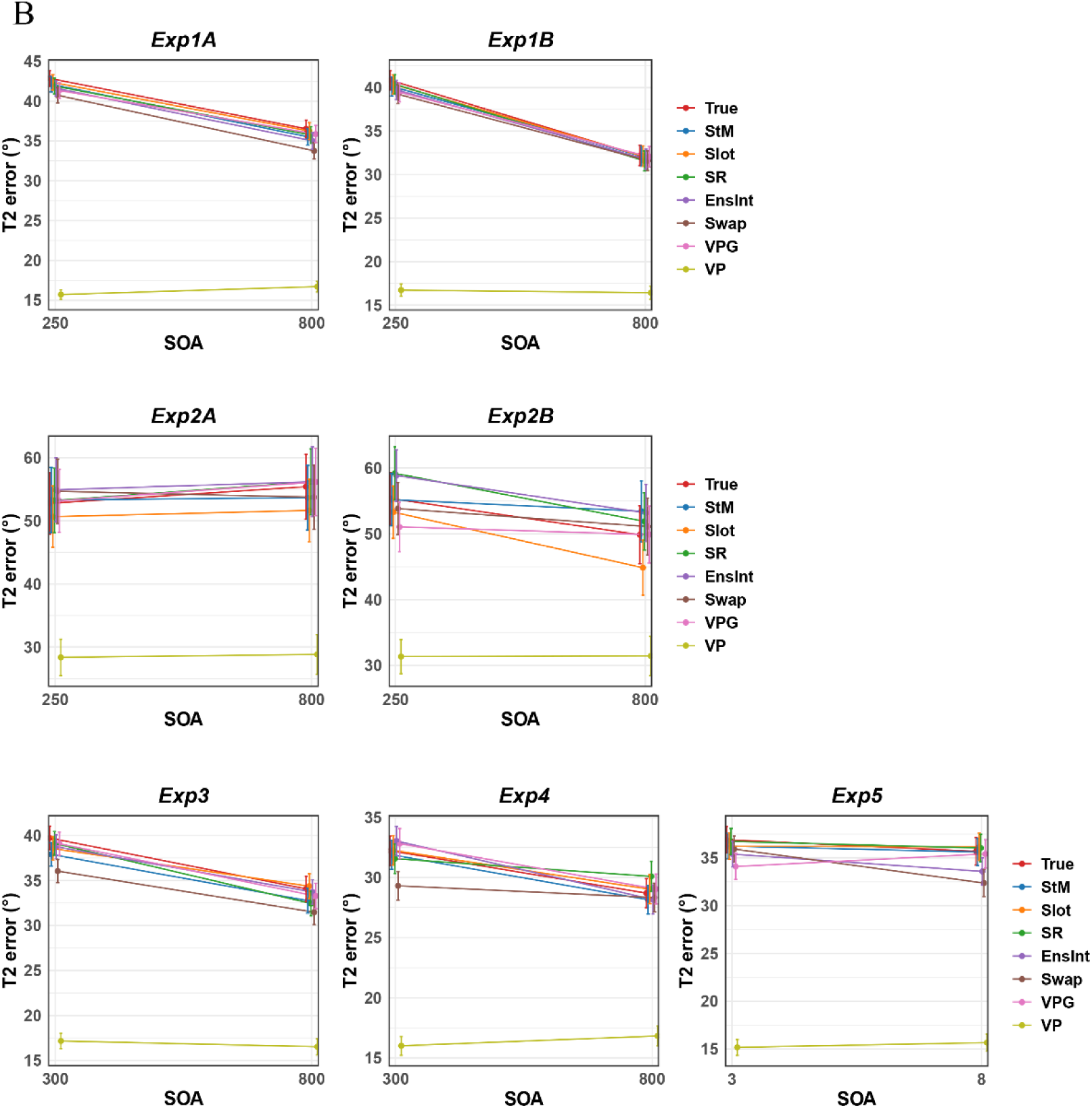
Comparison of model simulations and experimental data based on T2 accuracy. *Note.* Model simulation results using the data from Karabay et al. (2022). (**A**) Average T2 errors in the T1 correct condition, shown separately for each experiment. (**B**) Average T2 errors in the T1 incorrect condition. Exp = Experiment. True = experimentally observed data. See Method for model abbreviations.

### 3.4. Wang et al. data

#### 3.4.1. Model comparison

In the model comparison in the T1 correct condition, a standout characteristic of Wang et al.’s data, compared to prior studies, lies in the minimal discrepancies among the candidate models (see Figure 11).

**Figure 11.**
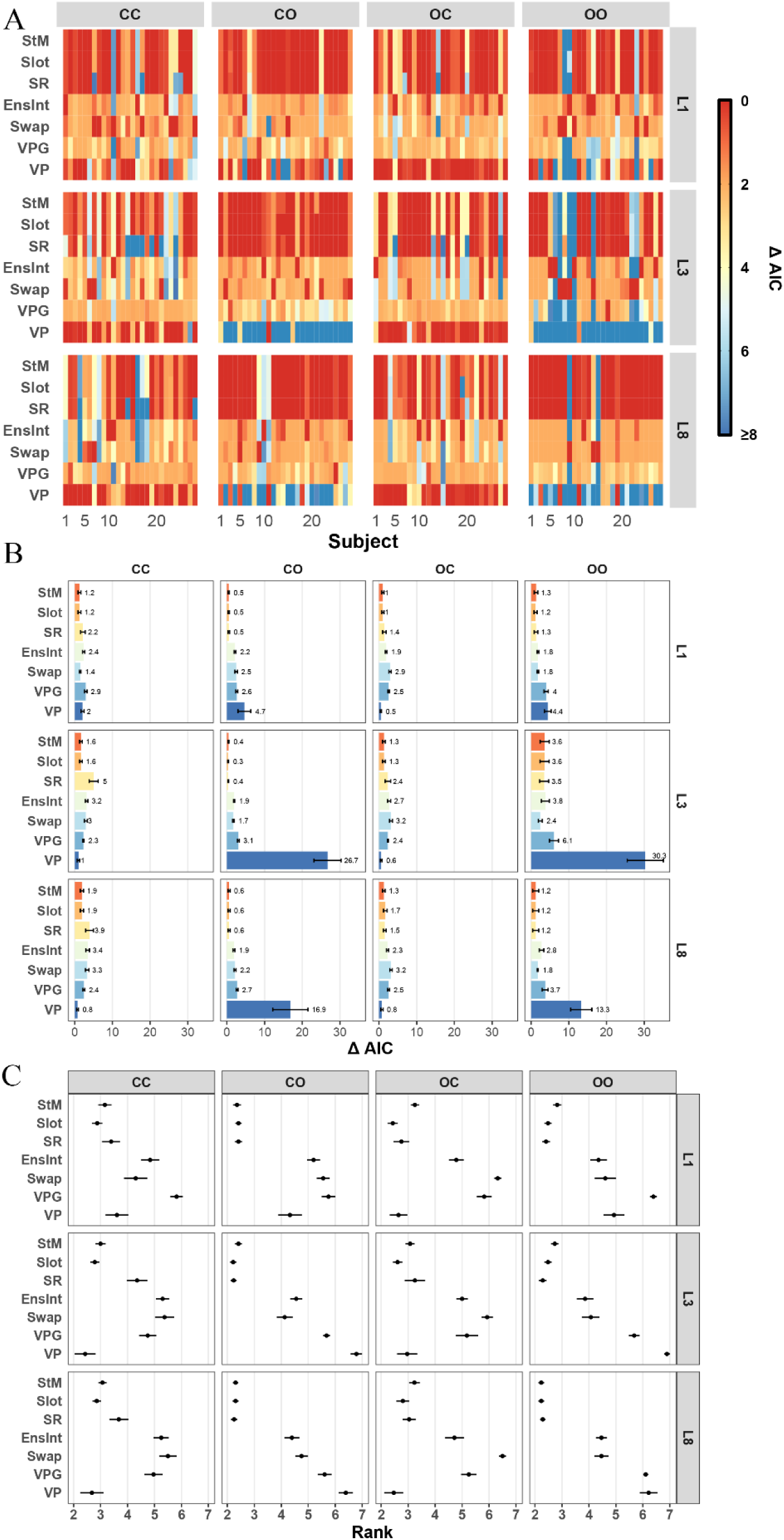
Model comparison results for T2 performance when T1 was correct (T2|T1). *Note.* (**A)** Model comparison at the subject level, using the data from Wang et al. The panels in each row present the results with the same lag but different target pairs; the panels in each column present the results with the same target pair but different lags. ‘CC’ means color-color target pair; ‘CO’ means color-orientation target pair; ‘OC’ means orientation-color target pair; ‘OO’ means orientation-orientation target pair. L1 = Lag 1; L3 = Lag 3; L8 = Lag 8. (**B**) Relative AIC values compared to that of the best-performing model, averaged across all subjects within each condition. (**C**) Average model ranking. Figure conventions follow Figure 2.

Analyzing at the subject level, we can group these models into three categories based on their performance (Figure 11A). First, the StM, Slot and SR models consistently demonstrated strong fitness across most subjects under different experimental conditions (lags and target pairs), as indicated by their comparatively lower or the lowest ΔAIC values. Notably, though, the SR model performed poorly for a subset of subjects when the color-color target pair was used. Second, the EnsInt, Swap and VPG models obtained moderate ΔAIC values across most subjects and various conditions, while the EnsInt and Swap models even best fitted several subjects at Lag 1 and 3, when both targets were colors or orientation gratings. Lastly, the VP model exhibited superior and stable performance when the second target was a color item – evident in the color-color and orientation-color target pair conditions. However, its fitness was worse when the second target featured orientation gratings. The VP model was competitive when T2 was color-based (CC/OC), but underperformed when T2 was orientation-based (OO/CO), mirroring the feature-dependent trend.

The narrow range of mean ΔAIC values for each experimental condition (from 0 to 30; refer to Figure 11B) further underscores the slight variances among models at the subject level. The VP model obviously underperformed when the second target was an orientation grating, as indicated by its highest mean ΔAIC value. Conversely, the VP model exhibited a comparable good fitness when the second target was a colored circle. Moreover, when two targets were both colors (CC), the SR model showed worse fitness than other models by presenting a higher mean ΔAIC value across all lags. Generally, the VPG, EnsInt and Swap models exhibited relatively inferior fitting across different experimental conditions, as indicated by their higher mean ΔAIC values. The average model ranking revealed a similar performance pattern (see Figure 11C). Overall, these feature-dependent model comparison results suggest that the performance of a model is influenced by the stimulus features employed in the experiments.

Figure 12 illustrates the assessment of these models’ fit to T2 data when T1 was incorrect. Because there were insufficient trials for each combination of lag and target pair, we confined our investigation to T2 performance at each lag. Thereby we can still provide an overview of the models’ performance in scenarios where only one item was successfully memorized into working memory. The overall pattern was similar to that seen when T1 was correct. For example, the VP model underperformed compared to the others. Notably, while the Swap model demonstrated superior fitness at Lag 1, these results must be interpreted with caution, due to the combined inclusion of both same- and different-feature target pairs.

**Figure 12.**
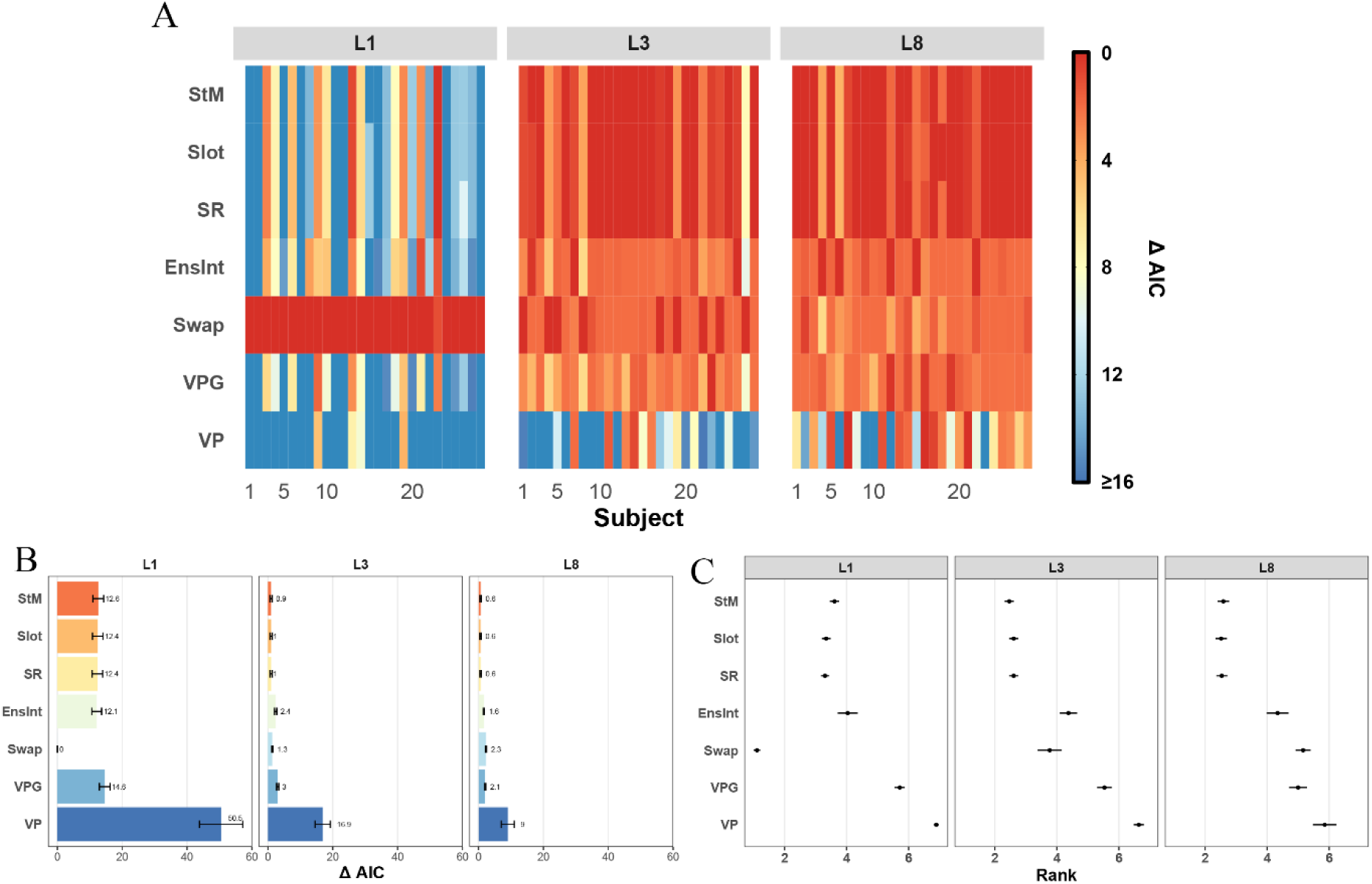
Model comparison results for T2 performance when T1 was incorrect. *Note.* (**A**) Model comparison at the subject level, using the data from Wang et al. L1 = Lag 1; L3 = Lag 3; L8 = Lag 8. (**B**) Relative AIC values compared to that of the best-performing model, averaged across all subjects at each specific lag. (**C**) Average model ranking at each lag. Figure conventions follow Figure 2.

#### 3.4.2. Model Simulation

The results of simulation analysis for the Wang et al. data are depicted in Figure 12. For the T1 correct condition (as shown in Figure 12A), the VPG model did not accurately represent the increased T2 error at Lag 3 when the second target was an orientation grating. Yet, when the second target was a color, the estimated T2 errors obtained from all models and the actual T2 errors clustered closely. It should be noted that there was no attentional blink detected in the experimental data at Lag 3, when the second target was a color. For the T1 incorrect condition (Figure 12B), a comparison between the simulated and actual data at each lag is possible. Here, the VPG model’s estimation of the T2 error was notably less accurate.

**Figure 12.**
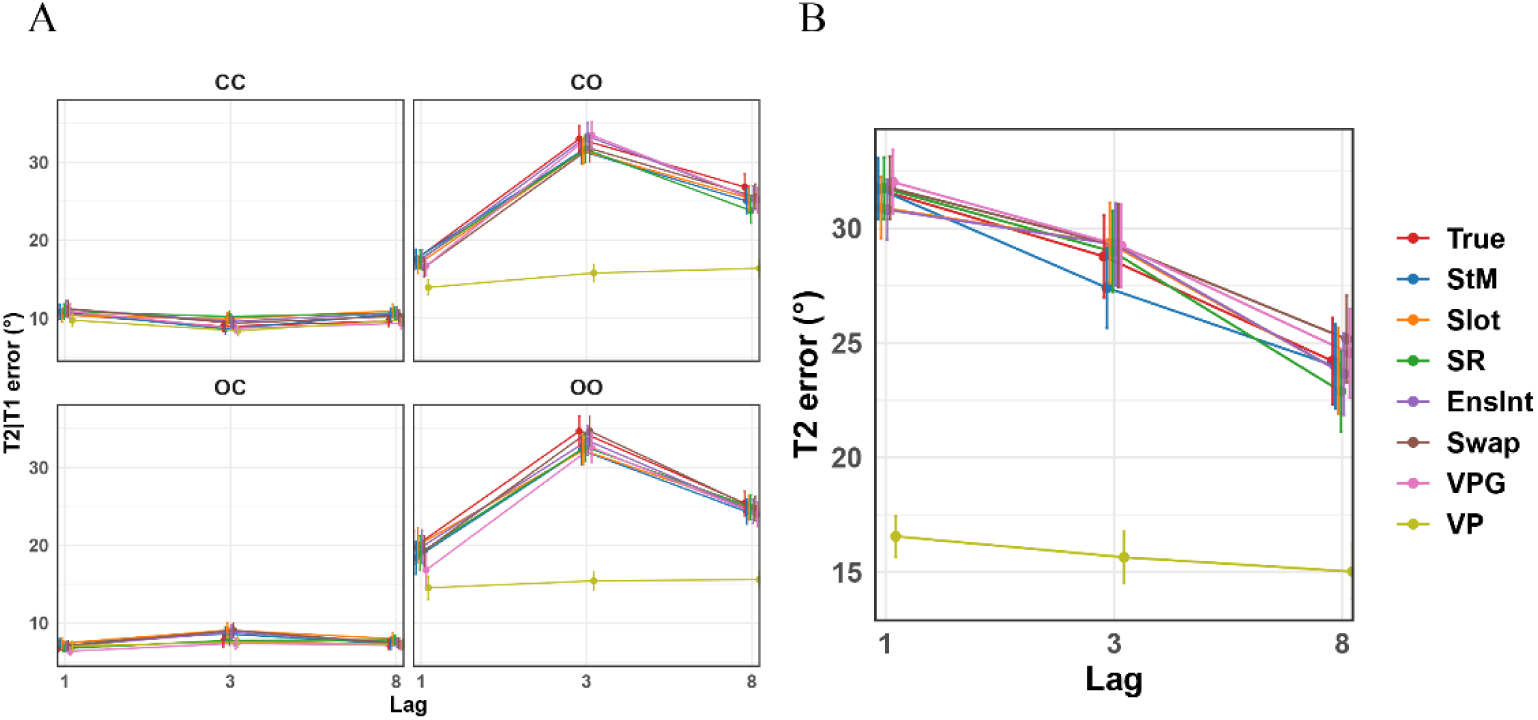
Comparison of model simulations and experimental data based on T2 accuracy. *Note.* Model simulation results using the data from Wang et al. (**A**) Simulation results in the T1 correct condition. Each panel presents the comparison of the mean T2 error in actual data and in data simulated from seven models, separately for each lag and each target pair condition. ‘CC’ means color-color target pair; ‘CO’ means color-orientation target pair; ‘OO’ means orientation-orientation target pair; ‘OC’ means orientation-color target pair. True = experimentally observed data. See Method for model abbreviations. (**B**) Simulation results in the T1 incorrect condition. The results were obtained across all target pairs within each specific lag.

## 4. Discussion

In the current study, we explored visual (WM) representations during AB tasks by systematically comparing a broader set of WM models across multiple datasets. Quantitative assessment of target representations in AB tasks by fitting WM models to continuous reproduction data has been done before, but previous studies have not reached a consensus on interpreting the error distribution of these WM representations for T2, partly due to a lack of systematic comparisons of these models in the AB domain and limited attention to models like the EnsInt and Swap models that consider potentially important nontarget reports. We described comparative fits of seven WM models to four sets of experimental data from three different laboratories. Specifically, we applied MLE for parameter estimation, which allowed us to assess the fitness of these models to the data and then compare them based on AIC scores. Additionally, we utilized the parameter values from each model to generate simulated data sets, further evaluating the accuracy of these models in describing the experimental data under various conditions.

Across datasets, we found that slot-family models most often win at short lags, supporting predominantly discrete consolidation failures during the AB. Yet, VP/VPG models improve with colors and longer lags, indicating that graded precision reductions emerge when timing/feature content affords partial consolidation. Overall, AB errors are thus mixed: They consist mostly of guesses at short lags, with more low-precision reports at longer lags or with color features. This reconciles prior mixed findings, and from a practical perspective suggests that one’s model choice should depend on the lag and feature dimensions being studied.

The three leading models, the StM, Slot and SR models, share a crucial assumption that only a limited number items can be encoded into WM. In these models’ error distributions, when items were not encoded into WM successfully, the corresponding errors follow a uniform distribution indicative of random responses. Our findings showed robust and consistent performance of these models across different data sets, and importantly, their simulations closely reproduced the ‘blink’ pattern in T2|T1 errors. This pattern of results supports existing WM encoding accounts for the AB phenomenon, where WM consolidation processes are disrupted during the blink period. Specifically, in rapid sequences of stimuli, T2 cannot be consolidated into WM until T1 is completely encoded (Bowman & Wyble, 2007; Chun & Potter, 1995; Jolicœur & Dell’Acqua, 1998; Wyble et al., 2009). These accounts suggest that WM consolidation is slow and capacity limited. In line with this idea, Ricker and Hardman (2017) provided a comprehensive analysis of the relationship between consolidation time and a stable WM representation. They explicitly associated the temporal dynamics of WM consolidation with the AB. They suggested that the duration of consolidation has a direct influence on the probability of a memory item being accessible during reporting.

In addition, the comparison results of the VP and VPG models further supported the importance of including a guess rate component in account for WM representation errors during the AB. First, the VP model generally underperformed at shorter lags (SOAs), indicating a failure to capture impaired T2 representations during blink period. At longer lags (SOAs), when there is sufficient time for both T1 and T2 consolidation, the VP model’s performance improved. Second, the VP model displayed enhanced performance under conditions of a weakened AB, such as in Experiment 2A and 2B of Karabay et al. (2022), as well as with color-color and orientation-color target pairs in the datasets of Wang et al. Most importantly, the VPG model, a variant of the VP model that incorporates a guess rate, showed better performance compared to the VP model. The simulation analysis unveiled several issues of the VP model: An underestimation of the response errors, and, more specifically, an underestimation of the response errors at shorter lags (SOAs) in relation to longer lags (SOAs), demonstrating a failure to accurately capture impaired memory consolidation during the AB. Contrastingly, the VPG model proved to be better, exhibiting a tendency to align with the empirical data across various conditions. These findings align with the results reported in Experiment 3 of Sy et al. (2021)., Our analysis thus suggests that WM errors during the AB are more effectively explained through guessing, as opposed to extremely low precision. This is in keeping with extant literature, which claims that T2 awareness can be discrete during the AB (Asplund et al., 2014; Sy et al., 2021; Wang et al., 2024). However, it is important to note that awareness during the AB can also be gradual, depending on the spatial extent of attention (Karabay et al., 2022).

Upon closer examination of the ‘slot’ models (the StM, Slot, and SR models), one might argue that the principal divergence lies in their predictions regarding the number of items that can be successfully encoded into WM, also referred to as set size. This is illustrated by the varying guess rates (*Pg*) for the Slot and SR models, which are determined based on distinct mathematical calculations that take into account the predicted number of available memory slots. To address this, we expanded our analysis to consider both T2|T1 performance, indicative of a set size of two, and T2 performance when T1 was incorrect, as a proxy of a set size of one. By doing so, we can gain insights into model fitness under different memory load conditions. Overall, the StM, Slot, and SR models exhibited similar performance across set sizes, and we could not reliably differentiate them for set sizes of one and two under most conditions. This lack of differentiation could nevertheless also be a consequence of the limited set sizes available in our analyzed data sets (one or two), which are presumably fewer than the number of available visual WM slots as suggested by Cowan (2001). In these analyses, we also noticed that T2 report performance was poorer in the T1 incorrect condition than in the T1 correct condition. We suggest that attentional resources might not be effectively focused in these trials overall, which may explain the poorer performance on T2. In this case, future AB research with set sizes pushing the WM limits will need to be undertaken (e.g., Akyürek et al., 2007) to further distinguish these models’ predictions and clarify the relationship between set size and WM consolidation during the AB.

Our study also investigated the performance of models that consider nontarget reports (the EnsInt and Swap models) in the AB domain. These models exhibited distinct performance patterns across different temporal intervals and feature dimensions. The EnsInt model did not show a clear pattern across datasets, making it difficult to definitively ascertain whether the EnsInt model’s fitness was impacted by different time intervals. By contrast, the Swap model’s performance appeared to be more influenced by the time interval between the two targets, s. For instance, the Swap model performed better at shorter lags, as evidenced in the model comparison results of Tang et al.’s datasets (see Figure 5 and 6), in the single-stream RSVP paradigm of Karabay et al.’s study (e.g., Experiment 3), and also in the same feature-dimension pairs (e.g., color-color and orientation-orientation) of Wang et al.’s data. These results were in accordance with previous research indicating that temporal target integration and target report order reversals occur frequently at Lag 1 (Akyürek et al., 2012; Karabay & Akyürek, 2019). This, in turn, supports the explanation for Lag 1 sparing (i.e., the lack of an AB at that lag) that is offered by the eSTST model of the AB (Wyble et al., 2009), which holds that both targets are consolidated together when they successively follow each other without intervening distractors. This is particularly relevant in AB tasks where both targets share a common feature-dimension, as the Swap model’s prediction of the T2 error distribution can be influenced by the first target. It might thus be reasonable to conclude that the failure of the Swap model at longer lags could be attributed to the weaker influence of T1 on T2 during memory consolidation. Moreover, when analyzing different set sizes, the Swap model displayed better fitness when the set size was one (T1 incorrect condition) compared to its performance when the set size was two (T1 correct condition), as noted in the datasets from Karabay et al. and Wang et al. This is likely due to the swap errors at short lags or SOAs, which result in better performance of the Swap model when T1 is incorrect. Overall, these findings showed that swap errors are present at shorter lags, indicating that T1 and T2 representations partially overlap during consolidation. This pattern implies that AB accounts can be extended by considering the possibility of misbinding between temporally adjacent targets.

## 5. Conclusion

In this study, we explored WM representations during the AB by evaluating seven WM models’ fitness against distinct data sets derived from AB tasks. Models positing a limited number of ‘storage slots’ in WM (StM, Slot, and SR) consistently outperformed other models. These results suggest that memory representations can be categorized into two distinct types: One representing a complete guess, which occurs when the item was not encoded into WM at all (due to a capacity limitation or processing limitation), and the other representing the precision with which information is encoded into visual WM. This dichotomy is implemented in the StM, Slot and SR models. Models accounting for long tails in error distributions further confirmed that guesses play a key role in explaining T2 representations during the AB (i.e., the contrast of VPG and VP models). Together, these results align with the theoretical AB models proposing that WM consolidation is disrupted, resulting in no conscious access to T2 during the AB (e.g., Asplund et al., 2014; Wyble et al., 2009).

Moreover, we identified that the lag between targets, the common feature-dimensions shared by targets, and the specific dimension of the target feature (color or orientation) played an important role in model performance. Task conditions thus affect the utility of different models, highlighting the need for careful consideration when selecting a model for a given experimental context. If a research question is centered around the short lags (lag 1-3), the EnsInt and Swap models should be preferred. Indeed, a model allowing both swaps and ensemble integrations should be considered for the short lags. However, models without target interactions should be preferred for later lags (lag 4-8; but see Tang et al., 2020). Future research will be needed to map out the role of attentional factors, such as target feature dimensions, in the WM consolidation process under time constraints. Such studies will help to clarify how overlapping T1-T2 representations and misbinding between targets contribute to memory errors, and to further guide model selection under different task conditions.

## Declarations

### Acknowledgements

We would like to thank Christopher L. Asplund for providing the original data sets from Asplund et al. (2014).

### Funding

Shuyao Wang was supported by the China Scholarship Council (CSC), grant 201906020179.

### Data availability

Data generated during the study (Wang et al. data sets), and all the analysis scripts are available on the Open Science Framework at: https://osf.io/zak5f/. The study was not preregistered. The data from Tang et al. (2020) and Karabay et al. (2022) are publicly available through their published articles. The Tang et al. (202) data can be accessed at: https://osf.io/f9g6h. The Karabay et al. (2022) data can be accessed at: https://osf.io/x5dru.

### Conflict of interest

The authors have no conflict of interest to disclose.

## Footnotes

1 Three subjects were removed from the original data set due to a lack of trials in the T1 incorrect condition. For consistency, we also excluded these subjects from all analyses of Asplund et al.’s data.

2 To verify the robustness of the fitting procedure, we conducted grid searches for the two VP parameters over wide ranges. Grid searches were performed on both the orientation and color datasets. Across all datasets, the best parameters from grid searches consistently fell at the lower boundary for the mode precision and showed very small values for the sd precision. This pattern closely aligned with the estimates obtained from MLE, indicating that the VP model’s poor performance is not due to optimization failure (e.g., the fitting procedure got stuck in a local minimum). Moreover, the model comparison results showed that the VPG model achieved better fits, which also suggest that the VP model’s poor performance reflects a mismatch between its predictions and the empirical data, rather than a parameter estimation issue.

3 For the data from Tang et al. (2022), we also conducted further analysis of the Ensemble Integration and Swap models, to investigate the potential interactions between T2 and its subsequent item of T2. Those analyses are presented in the Supplementary Materials (see Supplementary Materials B)

## Notes

The authors declare that they have no conflict of interest.

https://osf.io/zak5f/

### Competing Interest Statement

The authors have declared no competing interest.

### Summary of Updates

The Introduction, Discussion, and Conclusion sections have been expanded to provide a more robust integration of the empirical findings within the existing literature on attentional mechanisms.

