## Supplementary materials for "Evaluating working memory representations during the attentional blink: A comparative modeling approach"

**Supplementary Material**

**A. Further Model fitting and Model Comparison Results**

**1. Asplund et al. (2014).**

**Table S1**

*Model comparison results for T2 performance in the T1 correct condition (T2|T1).*

| Lag | Model | Mean AIC | Mean ΔAIC | Mean Rank | Mean Log-likelihood |
| --- | --- | --- | --- | --- | --- |
| 1 | StM | 1559.77 | 0.24 | 2.18 | -777.89 |
|  | Slot | 1559.77 | 0.24 | 2.18 | -777.89 |
|  | SR | 1559.77 | 0.24 | 2.12 | -777.89 |
|  | VPG | 1561.09 | 1.56 | 3.52 | -777.55 |
|  | VP | 1588.21 | 28.68 | 5 | -792.11 |
| 2 | StM | 1608.82 | 0.53 | 2.24 | -802.41 |
|  | Slot | 1608.82 | 0.53 | 2.24 | -802.41 |
|  | SR | 1608.82 | 0.53 | 2.24 | -802.41 |
|  | VPG | 1609.65 | 1.36 | 3.28 | -801.83 |
|  | VP | 1651.48 | 43.19 | 5 | -823.74 |
| 4 | StM | 1565.80 | 0.82 | 2.46 | -780.90 |
|  | Slot | 1565.80 | 0.82 | 2.4 | -780.90 |
|  | SR | 1565.80 | 0.82 | 2.46 | -780.90 |
|  | VPG | 1565.99 | 1.02 | 2.92 | -780.00 |
|  | VP | 1588.11 | 23.13 | 4.76 | -792.05 |
| 8 | StM | 1490.43 | 0.69 | 2.52 | -743.21 |
|  | Slot | 1490.43 | 0.69 | 2.52 | -743.21 |
|  | SR | 1490.43 | 0.69 | 2.52 | -743.21 |
|  | VPG | 1490.54 | 0.80 | 2.8 | -742.27 |
|  | VP | 1504.02 | 14.29 | 4.64 | -750.01 |

*Note.* AIC = Akaike information criterion. See *2.1. Model details* for the model abbreviation. Mean AIC values were obtained from averaging AIC values per model across all subjects, specifically for each lag. Mean ΔAIC values were obtained from averaging the relative AIC values compared to the best fitting model across all subjects, for each lag. For a detailed discussion of how these values were calculated, see *2.6. Model comparison* section of the main text.

**Table S2**

*Model comparison results for T2 performance in the T1 incorrect condition.*

| Lag | Model | Mean AIC | Mean ΔAIC | Mean Rank | Mean Log-likelihood |
| --- | --- | --- | --- | --- | --- |
| 1 | StM | 172.28 | 0.29 | 2.12 | -84.14 |
|  | Slot | 172.01 | 0.02 | 2.02 | -84.00 |
|  | SR | 172.01 | 0.02 | 1.98 | -84.00 |
|  | VPG | 174.39 | 2.40 | 4.36 | -84.20 |
|  | VP | 181.53 | 9.54 | 4.52 | -88.77 |
| 2 | StM | 153.55 | 0.10 | 2.12 | -74.78 |
|  | Slot | 153.45 | 0.00 | 1.96 | -74.73 |
|  | SR | 153.45 | 0.00 | 1.92 | -74.73 |
|  | VPG | 155.76 | 2.31 | 4.16 | -74.88 |
|  | VP | 166.70 | 13.25 | 4.84 | -81.35 |
| 4 | StM | 155.35 | 0.02 | 2.08 | -75.67 |
|  | Slot | 155.33 | 0.00 | 1.96 | -75.67 |
|  | SR | 155.33 | 0.00 | 1.96 | -75.67 |
|  | VPG | 157.49 | 2.16 | 4.24 | -75.75 |
|  | VP | 164.00 | 8.67 | 4.76 | -80.00 |
| 8 | StM | 171.64 | 0.15 | 2.2 | -83.82 |
|  | Slot | 171.50 | 0.01 | 2.04 | -83.75 |
|  | SR | 171.50 | 0.01 | 1.92 | -83.75 |
|  | VPG | 173.71 | 2.22 | 4.24 | -83.86 |
|  | VP | 182.49 | 11.00 | 4.6 | -89.24 |

**Table S3**

*Maximum-likelihood estimates of the parameters in all models in the T1 correct condition.*

|  |  | **Lag 1** | | **Lag 2** | | **Lag 4** | | **Lag 8** | |
| --- | --- | --- | --- | --- | --- | --- | --- | --- | --- |
| **Model** | **Parameter** | **Median** | ***SD*** | **Median** | ***SD*** | **Median** | ***SD*** | **Median** | ***SD*** |
| StM | *g* | 0.37 | 0.10 | 0.45 | 0.11 | 0.37 | 0.12 | 0.27 | 0.13 |
|  | *sd* | 21.21 | 3.49 | 20.69 | 4.34 | 20.76 | 4.37 | 19.23 | 4.06 |
| Slot | *capacity K* | 1.26 | 0.21 | 1.09 | 0.21 | 1.27 | 0.25 | 1.46 | 0.27 |
|  | *sd* | 21.21 | 3.49 | 20.69 | 4.34 | 20.76 | 4.37 | 19.23 | 4.06 |
| SR | *capacity K* | 1.26 | 0.21 | 1.09 | 0.21 | 1.27 | 0.25 | 1.46 | 0.27 |
|  | *sd* | 19.53 | 3.16 | 19.89 | 3.45 | 19.56 | 3.91 | 16.10 | 3.72 |
| VPG | *g* | 0.33 | 0.11 | 0.39 | 0.14 | 0.30 | 0.13 | 0.21 | 0.14 |
|  | *modePrecision* | 2.42×10^-3^ | 8.56×10^-4^ | 2.65×10^-3^ | 1.45×10^-3^ | 2.34×10^-3^ | 1.47×10^-3^ | 3.5×10^-3^ | 1.76×10^-3^ |
|  | *sdPrecision* | 1.20×10^-3^ | 5.5×10^-3^ | 1.62×10^-3^ | 5.96×10^-3^ | 3.79×10^-3^ | 5.03×10^-3^ | 3.81×10^-3^ | 4.63×10^-3^ |
| VP | *modePrecision* | 1×10^-3^ | 1.98×10^-11^ | 1×10^-3^ | 5.78×10^-12^ | 1×10^-3^ | 3.42×10^-4^ | 1×10^-3^ | 1.6×10^-3^ |
|  | *sd*Precision | 4.65×10^-3^ | 2.5×10^-3^ | 4.62×10^-3^ | 2.84×10^-3^ | 5.19×10^-3^ | 3.49×10^-3^ | 8.79×10^-3^ | 1.25×10^-2^ |

*Note.* See *2.1. Model details* of the main text for the model abbreviations.

**Table S4**

*Maximum-likelihood estimates of the parameters in all models in the T1 incorrect condition.*

|  |  | **Lag 1** | | **Lag 2** | | **Lag 4** | | **Lag 8** | |
| --- | --- | --- | --- | --- | --- | --- | --- | --- | --- |
| **Model** | **Parameter** | **Median** | ***SD*** | **Median** | ***SD*** | **Median** | ***SD*** | **Median** | ***SD*** |
| StM | *g* | 0.36 | 0.29 | 0.49 | 0.34 | 0.39 | 0.29 | 0.28 | 0.31 |
|  | *sd* | 25.75 | 33.79 | 21.18 | 140.81 | 22.35 | 24.61 | 28.46 | 21.84 |
| Slot | *capacity K* | 1.10 | 1.40 | 1.02 | 0.97 | 1.21 | 0.72 | 1.44 | 0.68 |
|  | *sd* | 21.06 | 31.48 | 18.24 | 29.12 | 22.30 | 25.57 | 19.58 | 22.05 |
| SR | *capacity K* | 1.14 | 0.78 | 1.03 | 1.22 | 1.21 | 0.86 | 1.44 | 0.92 |
|  | *sd* | 23.51 | 21.10 | 27.00 | 75.75 | 18.64 | 19.05 | 19.13 | 17.65 |
| VPG | *g* | 0.50 | 0.30 | 0.58 | 0.31 | 0.50 | 0.29 | 0.31 | 0.32 |
|  | *modePrecision* | 1.88×10^-3^ | 4.17 | 3.56×10^-3^ | 6.18 | 1.66×10^-3^ | 0.03 | 1.86×10^-3^ | 1.31 |
|  | *sdPrecision* | 1×10^-3^ | 33.04 | 1.06×10^-3^ | 27.69 | 1×10^-3^ | 20 | 1×10^-3^ | 20 |
| VP | *modePrecision* | 1×10^-3^ | 1.1×10^-3^ | 1×10^-3^ | 1.03×10^-2^ | 1×10^-3^ | 9.64×10^-4^ | 1×10^-3^ | 2.76×10^-3^ |
|  | *sd*Precision | 3.21×10^-3^ | 0.18 | 3.24×10^-3^ | 6.7×10^-3^ | 3.54×10^-3^ | 20 | 3.31×10^-3^ | 1.51×10^-2^ |

**2. Tang et al. (2020).**

**Table S5**

*Model comparison results for T2 performance in the T1 correct condition (T2|T1).*

| **Experiment** | **Lag** | **Model** | **Mean AIC** | **Mean ΔAIC** | **Mean**  **Rank** | **Mean**  **Log-likelihood** |
| --- | --- | --- | --- | --- | --- | --- |
| ***Exp1*** | **1** | StM | 318.28 | 9.92 | 3.34 | -157.14 |
|  |  | Slot | 318.23 | 9.88 | 3.07 | -157.12 |
|  |  | SR | 318.20 | 9.84 | 3.00 | -157.10 |
|  |  | EnsInt | 315.76 | 7.40 | 2.93 | -154.88 |
|  |  | Swap | 309.52 | 1.17 | 2.75 | -151.76 |
|  |  | VPG | 320.86 | 12.50 | 5.91 | -157.43 |
|  |  | VP | 353.49 | 45.13 | 7.00 | -174.74 |
|  | **2** | StM | 509.62 | 16.89 | 3.32 | -252.81 |
|  |  | Slot | 509.35 | 16.62 | 2.84 | -252.68 |
|  |  | SR | 509.34 | 16.60 | 2.93 | -252.67 |
|  |  | EnsInt | 506.71 | 13.97 | 3.25 | -250.35 |
|  |  | Swap | 494.16 | 1.42 | 2.93 | -244.08 |
|  |  | VPG | 512.04 | 19.30 | 5.73 | -253.02 |
|  |  | VP | 575.61 | 82.87 | 7.00 | -285.80 |
|  | **3** | StM | 510.92 | 12.29 | 3.14 | -253.46 |
|  |  | Slot | 510.72 | 12.09 | 2.89 | -253.36 |
|  |  | SR | 510.69 | 12.07 | 2.84 | -253.35 |
|  |  | EnsInt | 507.99 | 9.36 | 3.64 | -250.99 |
|  |  | Swap | 499.56 | 0.94 | 2.64 | -246.78 |
|  |  | VPG | 514.18 | 15.56 | 5.86 | -254.09 |
|  |  | VP | 565.16 | 66.53 | 7.00 | -280.58 |
|  | **5** | StM | 546.46 | 5.81 | 2.89 | -271.23 |
|  |  | Slot | 546.38 | 5.74 | 2.64 | -271.19 |
|  |  | SR | 546.38 | 5.74 | 2.66 | -271.19 |
|  |  | EnsInt | 545.27 | 4.63 | 4.30 | -269.64 |
|  |  | Swap | 541.91 | 1.27 | 3.48 | -267.96 |
|  |  | VPG | 549.03 | 8.39 | 5.55 | -271.52 |
|  |  | VP | 582.04 | 41.39 | 6.50 | -289.02 |
|  | **7** | StM | 498.98 | 2.64 | 2.61 | -247.49 |
|  |  | Slot | 498.90 | 2.56 | 2.50 | -247.45 |
|  |  | SR | 498.83 | 2.50 | 2.52 | -247.42 |
|  |  | EnsInt | 500.10 | 3.77 | 4.66 | -247.05 |
|  |  | Swap | 497.77 | 1.43 | 3.80 | -245.88 |
|  |  | VPG | 501.87 | 5.53 | 5.73 | -247.93 |
|  |  | VP | 525.78 | 29.45 | 6.18 | -260.89 |

**Table S5** (*continued*)

| **Experiment** | **Lag** | **Model** | **Mean AIC** | **Mean ΔAIC** | **Mean rank** | **Log-likelihood** |
| --- | --- | --- | --- | --- | --- | --- |
| ***Exp2*** | 3 | StM | 1356.37 | 3.87 | 2.59 | -676.19 |
|  |  | Slot | 1356.37 | 3.87 | 2.59 | -676.19 |
|  |  | SR | 1356.37 | 3.87 | 2.65 | -676.19 |
|  |  | EnsInt | 1356.47 | 3.97 | 4.13 | -675.23 |
|  |  | Swap | 1353.60 | 1.10 | 3.30 | -673.80 |
|  |  | VPG | 1360.03 | 7.53 | 5.78 | -677.02 |
|  |  | VP | 1437.18 | 84.68 | 6.96 | -716.59 |
|  | **7** | StM | 1306.75 | 1.23 | 2.46 | -651.38 |
|  |  | Slot | 1306.95 | 1.44 | 2.59 | -651.48 |
|  |  | SR | 1306.57 | 1.06 | 2.35 | -651.29 |
|  |  | EnsInt | 1308.29 | 2.77 | 4.70 | -651.14 |
|  |  | Swap | 1307.78 | 2.26 | 4.43 | -650.89 |
|  |  | VPG | 1311.27 | 5.75 | 5.61 | -652.63 |
|  |  | VP | 1340.81 | 35.29 | 5.87 | -668.40 |

**Table S6**

*Model comparison results for T2 performance in the T1 incorrect condition.*

| **Experiment** | **Lag** | **Model** | **Mean AIC** | **Mean ΔAIC** | **Mean**  **Ranking** | **Mean**  **Log-likelihood** |
| --- | --- | --- | --- | --- | --- | --- |
| ***Exp1*** | **1** | StM | 769.12 | 1.68 | 2.61 | -382.56 |
|  |  | Slot | 769.08 | 1.64 | 2.50 | -382.54 |
|  |  | SR | 769.06 | 1.63 | 2.43 | -382.53 |
|  |  | EnsInt | 770.26 | 2.82 | 4.43 | -382.13 |
|  |  | Swap | 768.50 | 1.06 | 3.39 | -381.25 |
|  |  | VPG | 771.90 | 4.46 | 5.73 | -382.95 |
|  |  | VP | 847.98 | 80.54 | 6.91 | -421.99 |
|  | **2** | StM | 587.21 | 2.77 | 2.93 | -291.60 |
|  |  | Slot | 587.21 | 2.77 | 2.82 | -291.60 |
|  |  | SR | 587.10 | 2.66 | 2.52 | -291.55 |
|  |  | EnsInt | 586.94 | 2.50 | 3.75 | -290.47 |
|  |  | Swap | 586.24 | 1.81 | 3.20 | -290.12 |
|  |  | VPG | 589.48 | 5.05 | 5.77 | -291.74 |
|  |  | VP | 661.21 | 76.77 | 7.00 | -328.60 |
|  | **3** | StM | 572.66 | 2.75 | 2.82 | -284.33 |
|  |  | Slot | 572.52 | 2.70 | 3.37 | -284.26 |
|  |  | SR | 572.25 | 2.42 | 2.55 | -284.12 |
|  |  | EnsInt | 572.73 | 2.82 | 3.13 | -283.37 |
|  |  | Swap | 572.23 | 2.32 | 2.41 | -283.11 |
|  |  | VPG | 575.15 | 5.23 | 2.89 | -284.57 |
|  |  | VP | 642.13 | 72.22 | 3.64 | -319.07 |
|  | **5** | StM | 521.61 | 1.09 | 3.57 | -258.81 |
|  |  | Slot | 521.55 | 1.02 | 3.77 | -258.77 |
|  |  | SR | 521.44 | 0.92 | 2.57 | -258.72 |
|  |  | EnsInt | 522.22 | 1.70 | 5.82 | -258.11 |
|  |  | Swap | 522.24 | 1.71 | 5.48 | -258.12 |
|  |  | VPG | 524.51 | 3.99 | 7.00 | -259.26 |
|  |  | VP | 573.37 | 52.84 | 7.00 | -284.68 |
|  | **7** | StM | 519.18 | 0.69 | 2.61 | -257.59 |
|  |  | Slot | 519.18 | 0.69 | 2.50 | -257.59 |
|  |  | SR | 519.18 | 0.69 | 2.30 | -257.59 |
|  |  | EnsInt | 520.26 | 1.77 | 3.89 | -257.13 |
|  |  | Swap | 520.49 | 2.00 | 4.11 | -257.25 |
|  |  | VPG | 522.52 | 4.03 | 5.91 | -258.26 |
|  |  | VP | 567.44 | 48.95 | 6.68 | -281.72 |
| ***Exp2*** | **3** | StM | 1267.25 | 8.34 | 2.45 | -631.62 |
|  |  | Slot | 1267.13 | 8.22 | 2.59 | -631.56 |
|  |  | SR | 1266.99 | 8.04 | 2.39 | -631.50 |
|  |  | EnsInt | 1265.45 | 6.53 | 2.59 | -629.72 |
|  |  | Swap | 1259.64 | 0.73 | 2.25 | -626.82 |
|  |  | VPG | 1270.05 | 11.14 | 2.48 | -632.02 |
|  |  | VP | 1410.50 | 151.59 | 4.05 | -703.25 |
|  | **7** | StM | 1073.56 | 0.82 | 4.17 | -534.78 |
|  |  | Slot | 1073.56 | 0.82 | 4.50 | -534.78 |
|  |  | SR | 1073.45 | 0.71 | 3.74 | -534.73 |
|  |  | EnsInt | 1074.63 | 1.88 | 5.86 | -534.31 |
|  |  | Swap | 1074.13 | 1.39 | 5.83 | -534.06 |
|  |  | VPG | 1077.55 | 4.81 | 6.50 | -535.77 |
|  |  | VP | 1151.23 | 78.49 | 6.61 | -573.61 |

**Table S7**

*Maximum-likelihood estimates of the parameters in all models in the T1 correct condition.*

|  |  | ***Exp1*** | | | | | | | | | | ***Exp2*** | | | |
| --- | --- | --- | --- | --- | --- | --- | --- | --- | --- | --- | --- | --- | --- | --- | --- |
|  |  | **Lag 1** | | **Lag 2** | | **Lag 3** | | **Lag 5** | | **Lag 7** | | **Lag 3** | | **Lag 7** | |
| **Model** | **Parameter** | **Median** | **SD** | **Median** | **SD** | **Median** | **SD** | **Median** | **SD** | **Median** | **SD** | **Median** | **SD** | **Median** | **SD** |
| StM | *g* | 0.65 | 0.41 | 0.83 | 0.44 | 0.73 | 0.43 | 0.48 | 0.38 | 0.19 | 0.38 | 0.31 | 0.34 | 0.16 | 0.28 |
|  | *sd* | 22.43 | 19.73 | 20.94 | 18.44 | 20.94 | 17.14 | 19.12 | 15.62 | 20.52 | 10.96 | 21.8 | 18.11 | 19.96 | 10.25 |
| Slot | *capacity K* | 0.56 | 1.47 | 0.33 | 1.36 | 0.5 | 1.53 | 1.02 | 0.94 | 1.53 | 1.2 | 1.38 | 1.47 | 1.67 | 1.40 |
|  | *sd* | 21.19 | 20.32 | 15.13 | 21.59 | 28.24 | 29.23 | 18.47 | 27.64 | 24.2 | 25 | 22.01 | 18.23 | 20.58 | 10.59 |
| SR | *capacity K* | 0.78 | 1.15 | 1.27 | 1.19 | 0.9 | 1.07 | 1.22 | 1.3 | 1.53 | 0.72 | 1.38 | 0.92 | 1.67 | 0.97 |
|  | *sd* | 27.03 | 50.06 | 35.59 | 73.62 | 23.43 | 50.35 | 21.27 | 40.96 | 20.99 | 55.96 | 26.15 | 39.25 | 16.43 | 40.47 |
| EnsInt | *g* | 0.26 | 0.37 | 2.47×10^-3^ | 0.43 | 0.39 | 0.38 | 0.18 | 0.37 | 0.15 | 0.28 | 0.29 | 0.3 | 0.13 | 0.25 |
|  | *sd* | 31.09 | 47.81 | 35.49 | 17.9 | 28.27 | 13.82 | 26.19 | 21.70 | 20.27 | 19.48 | 33.25 | 16.18 | 19.96 | 13.98 |
|  | *samples* | 4.43×10^-4^ | 4.59×10^12^ | 2.16×10^-5^ | 1.24×10^12^ | 0.22 | 7.04×10^12^ | 0.94 | 5.87×10^12^ | 10.28 | 3.78×10^12^ | 56.09 | 6.01×10^12^ | 8.23×10^12^ | 7.43×10^12^ |
| Swap | *g* | 0.08 | 0.28 | 0.07 | 0.27 | 0.09 | 0.36 | 0.15 | 0.23 | 0.05 | 0.23 | 0.04 | 0.25 | 0.12 | 0.23 |
|  | *B* | 0.41 | 0.4 | 0.47 | 0.37 | 0.15 | 0.38 | 0.04 | 0.37 | 1.97×10^-5^ | 0.35 | 0.05 | 0.27 | 6.88×10^-15^ | 0.22 |
|  | *sd* | 25.76 | 16.15 | 24.44 | 52.79 | 26.73 | 28.54 | 23.53 | 48.27 | 24.55 | 11.94 | 32.20 | 16.82 | 19.96 | 10.78 |
| VPG | *g* | 0.92 | 0.2 | 0.91 | 0.16 | 0.91 | 0.25 | 0.69 | 0.36 | 0.5 | 0.33 | 0.69 | 0.3 | 0.33 | 0.32 |
|  | *mode-*  *Precision* | 5.58×10^-4^ | 0.7 | 5.15×10^-4^ | 0.42 | 5.15×10^-4^ | 0.03 | 5.08×10^-4^ | 5.63×10^-4^ | 5×10^-4^ | 2.57×10^-3^ | 5×10^-4^ | 6.82×10^-4^ | 5×10^-4^ | 1.61×10^-4^ |
|  | *sdPrecision* | 5×10^-4^ | 10.66 | 7.89×10^-4^ | 10.66 | 6.76×10^-4^ | 14.71 | 5.38×10^-4^ | 0.05 | 6.99×10^-4^ | 10.66 | 5×10^-4^ | 0.01 | 5×10^-4^ | 3.6×10^-4^ |
| VP | *mode-*  *Precision* | 5×10^-4^ | 1.49×10^-11^ | 5×10^-4^ | 7.56×10^-12^ | 5×10^-4^ | 1.1×10^-11^ | 5×10^-4^ | 8.48×10^-5^ | 5×10^-4^ | 1.36×10^-4^ | 5×10^-4^ | 4.97×10^-12^ | 5×10^-4^ | 1.84×10^-4^ |
|  | *sd*Precision | 1.56×10^-3^ | 3.05×10^-4^ | 1.64×10^-3^ | 1.89×10^-4^ | 1.57×10^-3^ | 2.35×10^-4^ | 1.5×10^-3^ | 3.37×10^-4^ | 1.4×10^-3^ | 3.3×10^-3^ | 1.5×10^-3^ | 2.38×10^-4^ | 1.22×10^-3^ | 3.82×10^-4^ |

**Table S8**

*Maximum-likelihood estimates of the parameters in all models in the T1 incorrect condition.*

|  |  | ***Exp1*** | | | | | | | | | | ***Exp2*** | | | |
| --- | --- | --- | --- | --- | --- | --- | --- | --- | --- | --- | --- | --- | --- | --- | --- |
|  |  | **Lag 1** | | **Lag 2** | | **Lag 3** | | **Lag 5** | | **Lag 7** | | **Lag 3** | | **Lag 7** | |
| **Model** | **Parameter** | **Median** | **SD** | **Median** | **SD** | **Median** | **SD** | **Median** | **SD** | **Median** | **SD** | **Median** | **SD** | **Median** | **SD** |
| StM | *g* | 0.59 | 0.38 | 0.81 | 0.38 | 0.46 | 0.40 | 0.59 | 0.37 | 0.24 | 0.37 | 0.65 | 0.41 | 0.42 | 0.34 |
|  | *sd* | 18.36 | 14.81 | 15.91 | 21.55 | 22.47 | 23.34 | 17.07 | 17.2 | 22.81 | 45.88 | 21.72 | 18.4 | 20.61 | 9.13 |
| Slot | *capacity K* | 0.81 | 1.53 | 0.39 | 0.89 | 0.76 | 1.03 | 0.83 | 1.69 | 1.4 | 0.97 | 0.56 | 0.79 | 1.16 | 0.98 |
|  | *sd* | 18.86 | 17.14 | 18 | 22.96 | 21.27 | 21.15 | 17.07 | 18.14 | 31.77 | 15.44 | 24.22 | 27.62 | 20.61 | 9.52 |
| SR | *capacity K* | 0.84 | 1.01 | 0.68 | 0.92 | 1.09 | 0.95 | 0.83 | 1.13 | 1.45 | 1.18 | 0.56 | 1.22 | 1.16 | 0.83 |
|  | *sd* | 24.62 | 93.78 | 23.82 | 78.58 | 26.17 | 62.94 | 19.91 | 12.35 | 24.84 | 64.54 | 19.48 | 16.13 | 19.71 | 10.86 |
| EnsInt | *g* | 0.43 | 0.39 | 0.76 | 0.37 | 0.46 | 0.40 | 0.59 | 0.37 | 0.19 | 0.37 | 0.17 | 0.41 | 0.32 | 0.3 |
|  | *sd* | 25.74 | 20.19 | 13.38 | 18.01 | 20.75 | 23.22 | 17.07 | 16.73 | 26.72 | 23.65 | 25.13 | 19.88 | 21.06 | 16.06 |
|  | *samples* | 0.94 | 2.29×10^12^ | 0.01 | 9.68×10^11^ | 0.04 | 3.5×10^12^ | 7.48 | 2.47×10^12^ | 17.12 | 1.15×10^13^ | 0.64 | 3.68×10^12^ | 6.25 | 4.03×10^12^ |
| Swap | *g* | 3.26×10^-3^ | 0.34 | 0.42 | 0.38 | 0.33 | 0.31 | 0.38 | 0.34 | 0.36 | 0.37 | 0.11 | 0.32 | 0.35 | 0.29 |
|  | *B* | 0.07 | 0.37 | 0.14 | 0.32 | 0.06 | 0.32 | 0.07 | 0.15 | 1.33×10^-10^ | 0.06 | 0.2 | 0.29 | 0.03 | 0.09 |
|  | *sd* | 26.65 | 17.37 | 23.86 | 47.13 | 41.37 | 43.39 | 22.98 | 47.21 | 21.16 | 17.82 | 26.52 | 15.56 | 20.61 | 8.81 |
| VPG | *g* | 0.82 | 0.29 | 0.85 | 0.17 | 0.9 | 0.24 | 0.78 | 0.28 | 0.59 | 0.31 | 0.79 | 0.27 | 0.62 | 0.31 |
|  | *mode-*  *Precision* | 5×10^-4^ | 1.04×10^-3^ | 5.41×10^-4^ | 6.54×10^-4^ | 5.07×10^-4^ | 2.02×10^-3^ | 5.07×10^-4^ | 0.02 | 5×10^-4^ | 2.58×10^-4^ | 5.02×10^-4^ | 0.33 | 5×10^-4^ | 1.01×10^-3^ |
|  | *sdPrecision* | 5×10^-4^ | 2.68×10^-3^ | 5.01×10^-4^ | 10.7 | 5×10^-4^ | 6.88×10^-3^ | 5.08×10^-4^ | 17.56 | 5.1×10^-4^ | 0.02 | 5.47×10^-4^ | 17.22 | 5×10^-4^ | 3.33×10^-3^ |
| VP | *mode-*  *Precision* | 5×10^-4^ | 1.33×10^-11^ | 5×10^-4^ | 1.48×10^-11^ | 5×10^-4^ | 2.81×10^-11^ | 5×10^-4^ | 1.38×10^-11^ | 5×10^-4^ | 4.28×10^-5^ | 5×10^-4^ | 4.14×10^-12^ | 5×10^-4^ | 1.57×10^-4^ |
|  | *sdPrecision* | 1.56×10^-3^ | 2.16×10^-4^ | 1.65×10^-3^ | 1.63×10^-4^ | 1.56×10^-3^ | 3.23×10^-3^ | 1.54×10^-3^ | 2.17×10^-3^ | 1.42×10^-3^ | 3.09×10^-4^ | 1.61×10^-3^ | 3.03×10^-4^ | 1.41×10^-3^ | 0.01 |

**3. Karabay et al. (2022).**

**Table S9**

*Model comparison results for T2 performance in the T1 correct condition (T2|T1).*

|  |  | **Short SOA** | | | | **Long SOA** | | | |
| --- | --- | --- | --- | --- | --- | --- | --- | --- | --- |
| **Experiment** | **Model** | **Mean AIC** | **Mean ΔAIC** | **Mean Rank** | **Mean**  **Log-likelihood** | **Mean AIC** | **Mean ΔAIC** | **Mean Rank** | **Mean**  **Log-likelihood** |
| ***Exp1A*** | StM | 2174.28 | 0.96 | 2.26 | -1085.14 | 2269.40 | 0.51 | 2.20 | -1132.70 |
|  | Slot | 2174.28 | 0.96 | 2.13 | -1085.14 | 2269.40 | 0.51 | 2.20 | -1132.70 |
|  | SR | 2174.28 | 0.96 | 2.13 | -1085.14 | 2269.40 | 0.51 | 2.26 | -1132.70 |
|  | EnsInt | 2176.35 | 3.03 | 4.83 | -1085.17 | 2271.39 | 2.50 | 4.80 | -1132.69 |
|  | Swap | 2175.36 | 2.05 | 4.70 | -1084.68 | 2271.37 | 2.48 | 4.89 | -1132.68 |
|  | VPG | 2177.86 | 4.54 | 4.96 | -1085.93 | 2272.39 | 3.50 | 5.00 | -1133.19 |
|  | VP | 2302.45 | 129.13 | 7.00 | -1149.22 | 2329.82 | 60.93 | 6.65 | -1162.91 |
| ***Exp1B*** | StM | 2275.15 | 0.09 | 2.14 | -1135.57 | 2305.73 | 0.69 | 2.33 | -1150.86 |
|  | Slot | 2275.14 | 0.08 | 2.07 | -1135.57 | 2305.73 | 0.69 | 2.33 | -1150.86 |
|  | SR | 2275.14 | 0.08 | 2.07 | -1135.57 | 2305.73 | 0.69 | 2.33 | -1150.86 |
|  | EnsInt | 2278.87 | 3.81 | 4.93 | -1136.43 | 2307.72 | 2.68 | 5.07 | -1150.86 |
|  | Swap | 2277.22 | 2.16 | 4.83 | -1135.61 | 2307.70 | 2.66 | 5.36 | -1150.85 |
|  | VPG | 2278.69 | 3.63 | 4.95 | -1136.34 | 2307.80 | 2.76 | 4.52 | -1150.90 |
|  | VP | 2406.58 | 131.52 | 7.00 | -1201.29 | 2346.17 | 41.13 | 6.05 | -1171.08 |
| ***Exp2A*** | StM | 1826.96 | 6.94 | 3.65 | -911.48 | 1851.74 | 8.50 | 3.88 | -923.87 |
|  | Slot | 1826.97 | 6.94 | 3.71 | -911.48 | 1851.74 | 8.50 | 3.88 | -923.87 |
|  | SR | 1826.97 | 6.94 | 3.83 | -911.48 | 1851.74 | 8.50 | 3.88 | -923.87 |
|  | EnsInt | 1826.64 | 6.62 | 4.17 | -910.32 | 1851.78 | 8.55 | 5.00 | -922.89 |
|  | Swap | 1828.46 | 8.43 | 5.98 | -911.23 | 1853.36 | 10.13 | 6.35 | -923.68 |
|  | VPG | 1821.91 | 1.89 | 3.46 | -907.96 | 1844.86 | 1.63 | 2.73 | -919.43 |
|  | VP | 1824.77 | 4.75 | 3.19 | -910.39 | 1846.12 | 2.88 | 2.27 | -921.06 |
| ***Exp2B*** | StM | 2386.28 | 3.58 | 3.06 | -1191.14 | 2339.42 | 5.06 | 3.39 | -1167.71 |
|  | Slot | 2387.35 | 4.65 | 3.28 | -1191.67 | 2339.54 | 5.19 | 3.61 | -1167.77 |
|  | SR | 2386.33 | 3.64 | 3.11 | -1191.16 | 2339.42 | 5.06 | 3.44 | -1167.71 |
|  | EnsInt | 2388.24 | 5.55 | 5.70 | -1191.12 | 2340.95 | 6.60 | 5.61 | -1167.47 |
|  | Swap | 2388.26 | 5.57 | 5.74 | -1191.13 | 2341.08 | 6.73 | 6.13 | -1167.54 |
|  | VPG | 2384.27 | 1.57 | 2.93 | -1189.13 | 2335.63 | 1.27 | 2.89 | -1164.81 |
|  | VP | 2389.49 | 6.80 | 4.19 | -1192.74 | 2336.28 | 1.92 | 2.93 | -1166.14 |
| ***Exp3*** | StM | 1904.31 | 10.46 | 2.35 | -950.15 | 1861.28 | 0.81 | 2.31 | -928.64 |
|  | Slot | 1904.31 | 10.46 | 2.42 | -950.15 | 1861.36 | 0.89 | 2.35 | -928.68 |
|  | SR | 1904.31 | 10.46 | 2.35 | -950.15 | 1861.83 | 1.36 | 2.71 | -928.91 |
|  | EnsInt | 1903.19 | 9.34 | 4.35 | -948.59 | 1863.22 | 2.76 | 4.79 | -928.61 |
|  | Swap | 1895.25 | 1.41 | 3.90 | -944.63 | 1862.67 | 2.21 | 4.50 | -928.34 |
|  | VPG | 1908.80 | 14.95 | 5.75 | -951.40 | 1865.51 | 5.05 | 5.50 | -929.76 |
|  | VP | 2030.28 | 136.43 | 6.88 | -1013.14 | 1881.33 | 20.86 | 5.83 | -938.66 |
| ***Exp4*** | StM | 1888.70 | 0.84 | 2.19 | -942.35 | 1864.14 | 1.39 | 2.38 | -930.07 |
|  | Slot | 1888.70 | 0.84 | 2.31 | -942.35 | 1864.46 | 1.70 | 2.67 | -930.23 |
|  | SR | 1888.70 | 0.84 | 2.25 | -942.35 | 1864.14 | 1.39 | 2.38 | -930.07 |
|  | EnsInt | 1889.64 | 1.78 | 4.29 | -941.82 | 1865.36 | 2.60 | 4.60 | -929.68 |
|  | Swap | 1890.29 | 2.44 | 4.50 | -942.15 | 1865.66 | 2.90 | 4.77 | -929.83 |
|  | VPG | 1891.88 | 4.02 | 5.46 | -942.94 | 1866.86 | 4.10 | 4.83 | -930.43 |
|  | VP | 1959.16 | 71.30 | 7.00 | -977.58 | 1891.04 | 28.28 | 6.38 | -943.52 |
| ***Exp5*** | StM | 1442.75 | 0.55 | 2.24 | -719.38 | 1458.46 | 0.06 | 2.07 | -727.23 |
|  | Slot | 1442.87 | 0.66 | 2.28 | -719.43 | 1458.55 | 0.14 | 2.22 | -727.27 |
|  | SR | 1442.75 | 0.55 | 2.13 | -719.38 | 1458.46 | 0.06 | 2.07 | -727.23 |
|  | EnsInt | 1444.21 | 2.01 | 4.30 | -719.10 | 1460.39 | 1.99 | 4.89 | -727.20 |
|  | Swap | 1444.63 | 2.43 | 4.83 | -719.32 | 1460.46 | 2.05 | 5.24 | -727.23 |
|  | VPG | 1446.36 | 4.16 | 5.48 | -720.18 | 1460.80 | 2.40 | 5.13 | -727.40 |
|  | VP | 1511.01 | 68.81 | 6.74 | -753.51 | 1495.78 | 37.38 | 6.39 | -745.89 |

*Note.* SOA = stimulus onset asynchrony.

**Table S10**

*Model comparison results for T2 performance in the T1 incorrect condition.*

|  |  | **Short SOA** | | | | **Long SOA** | | | |
| --- | --- | --- | --- | --- | --- | --- | --- | --- | --- |
| **Experiment** | **Model** | **Mean AIC** | **Mean ΔAIC** | **Mean Rank** | **Mean**  **Log-likelihood** | **Mean AIC** | **Mean ΔAIC** | **Mean Rank** | **Mean**  **Log-likelihood** |
| ***Exp1A*** | StM | 1493.33 | 1.57 | 2.54 | -744.66 | 1282.36 | 1.71 | 2.67 | -639.18 |
|  | Slot | 1493.32 | 1.56 | 2.54 | -744.66 | 1282.15 | 1.51 | 2.50 | -639.08 |
|  | SR | 1493.27 | 1.52 | 2.48 | -744.64 | 1282.15 | 1.51 | 2.43 | -639.08 |
|  | EnsInt | 1495.01 | 3.25 | 4.87 | -744.50 | 1283.03 | 2.38 | 4.48 | -638.51 |
|  | Swap | 1492.98 | 1.22 | 3.13 | -743.49 | 1282.22 | 1.57 | 3.83 | -638.11 |
|  | VPG | 1495.30 | 3.54 | 5.43 | -744.65 | 1284.42 | 3.77 | 5.22 | -639.21 |
|  | VP | 1685.43 | 193.67 | 7.00 | -840.71 | 1392.00 | 111.35 | 6.87 | -694.00 |
| ***Exp1B*** | StM | 1275.35 | 1.93 | 2.67 | -635.68 | 1086.22 | 0.71 | 2.21 | -541.11 |
|  | Slot | 1275.35 | 1.93 | 2.67 | -635.68 | 1086.23 | 0.72 | 2.29 | -541.12 |
|  | SR | 1275.32 | 1.89 | 2.52 | -635.66 | 1086.22 | 0.71 | 2.21 | -541.11 |
|  | EnsInt | 1275.96 | 2.54 | 4.19 | -634.98 | 1088.10 | 2.59 | 5.07 | -541.05 |
|  | Swap | 1275.02 | 1.59 | 3.33 | -634.51 | 1086.99 | 1.48 | 4.17 | -540.49 |
|  | VPG | 1277.92 | 4.49 | 5.62 | -635.96 | 1088.80 | 3.29 | 5.33 | -541.40 |
|  | VP | 1423.83 | 150.40 | 7.00 | -709.91 | 1151.80 | 66.29 | 6.71 | -573.90 |
| ***Exp2A*** | StM | 186.56 | 2.81 | 2.75 | -91.28 | 169.49 | 2.05 | 2.83 | -82.75 |
|  | Slot | 185.61 | 1.86 | 2.54 | -90.80 | 169.49 | 2.05 | 2.60 | -82.75 |
|  | SR | 185.61 | 2.76 | 2.56 | -90.80 | 169.49 | 2.05 | 2.54 | -82.75 |
|  | EnsInt | 187.22 | 3.47 | 4.10 | -90.61 | 169.41 | 1.97 | 3.90 | -81.70 |
|  | Swap | 186.49 | 2.74 | 4.13 | -90.25 | 169.86 | 2.42 | 4.52 | -81.93 |
|  | VPG | 188.71 | 4.95 | 5.92 | -91.35 | 171.48 | 4.04 | 6.08 | -82.74 |
|  | VP | 199.17 | 15.42 | 6.00 | -97.58 | 180.99 | 13.55 | 5.54 | -88.49 |
| ***Exp2B*** | StM | 276.10 | 5.65 | 2.83 | -136.05 | 204.01 | 2.70 | 2.87 | -100.01 |
|  | Slot | 276.10 | 5.51 | 2.78 | -136.05 | 203.81 | 2.70 | 2.87 | -99.90 |
|  | SR | 276.10 | 5.70 | 2.98 | -136.05 | 204.01 | 2.50 | 2.67 | -100.01 |
|  | EnsInt | 276.46 | 6.00 | 4.41 | -135.23 | 203.74 | 2.43 | 4.06 | -98.87 |
|  | Swap | 271.14 | 0.69 | 2.44 | -132.57 | 203.24 | 1.93 | 3.50 | -98.62 |
|  | VPG | 278.23 | 7.78 | 6.11 | -136.11 | 206.12 | 4.81 | 6.15 | -100.06 |
|  | VP | 288.74 | 18.29 | 6.44 | -142.37 | 211.83 | 10.52 | 5.89 | -103.92 |
| ***Exp3*** | StM | 789.38 | 0.64 | 2.31 | -392.69 | 673.82 | 1.08 | 2.31 | -334.91 |
|  | Slot | 789.38 | 0.64 | 2.31 | -392.69 | 673.64 | 0.90 | 2.17 | -334.82 |
|  | SR | 789.38 | 0.64 | 2.25 | -392.69 | 673.64 | 0.90 | 2.06 | -334.82 |
|  | EnsInt | 791.22 | 2.48 | 4.65 | -392.61 | 674.61 | 1.87 | 4.52 | -334.31 |
|  | Swap | 790.13 | 1.39 | 3.77 | -392.07 | 675.54 | 2.80 | 4.73 | -334.77 |
|  | VPG | 792.02 | 3.27 | 5.79 | -393.01 | 676.47 | 3.73 | 5.92 | -335.23 |
|  | VP | 875.99 | 87.25 | 6.92 | -436.00 | 726.31 | 53.56 | 6.29 | -361.15 |
| ***Exp4*** | StM | 856.76 | 1.06 | 2.42 | -426.38 | 802.15 | 1.61 | 2.31 | -399.08 |
|  | Slot | 856.76 | 1.06 | 2.54 | -426.38 | 802.15 | 1.61 | 2.38 | -399.08 |
|  | SR | 856.75 | 1.05 | 2.38 | -426.37 | 802.15 | 1.61 | 2.44 | -399.08 |
|  | EnsInt | 858.20 | 2.51 | 4.77 | -426.10 | 804.06 | 3.52 | 4.98 | -399.03 |
|  | Swap | 856.90 | 1.20 | 3.40 | -425.45 | 801.62 | 1.08 | 3.35 | -397.81 |
|  | VPG | 859.53 | 3.84 | 5.50 | -426.77 | 805.14 | 4.60 | 5.58 | -399.57 |
|  | VP | 908.58 | 52.88 | 7.00 | -452.29 | 834.36 | 33.82 | 6.96 | -415.18 |
| ***Exp5*** | StM | 740.58 | 2.78 | 2.35 | -368.29 | 693.13 | 4.45 | 2.74 | -344.56 |
|  | Slot | 740.53 | 2.78 | 2.37 | -368.26 | 692.98 | 4.30 | 2.54 | -344.49 |
|  | SR | 740.57 | 2.72 | 2.15 | -368.28 | 692.91 | 4.23 | 2.37 | -344.45 |
|  | EnsInt | 741.38 | 3.58 | 4.63 | -367.69 | 692.84 | 4.16 | 4.00 | -343.42 |
|  | Swap | 739.43 | 1.63 | 4.37 | -366.71 | 691.13 | 2.45 | 4.26 | -342.56 |
|  | VPG | 742.56 | 4.76 | 5.39 | -368.28 | 695.19 | 6.51 | 5.52 | -344.59 |
|  | VP | 810.13 | 72.33 | 6.74 | -403.06 | 751.32 | 62.64 | 6.57 | -373.66 |

**Table S11**

*Maximum-likelihood estimates of the parameters in all models in the T1 correct condition.*

|  |  | ***Exp1A*** | | | | ***Exp1B*** | | | | ***Exp2A*** | | | | ***Exp2B*** | | | |
| --- | --- | --- | --- | --- | --- | --- | --- | --- | --- | --- | --- | --- | --- | --- | --- | --- | --- |
|  |  | **Short SOA** | | **Long SOA** | | **Short SOA** | | **Long SOA** | | **Short SOA** | | **Long SOA** | | **Short SOA** | | **Long SOA** | |
| **Model** | **Parameter** | **Median** | **SD** | **Median** | **SD** | **Median** | **SD** | **Median** | **SD** | **Median** | **SD** | **Median** | **SD** | **Median** | **SD** | **Median** | **SD** |
| **StM** | *g* | 0.36 | 0.3 | 0.31 | 0.18 | 0.56 | 0.25 | 0.35 | 0.18 | 0.06 | 0.11 | 0.06 | 0.10 | 0.09 | 0.11 | 0.06 | 0.05 |
|  | *sd* | 22.53 | 16.78 | 18.25 | 6.97 | 18.16 | 9.57 | 15.17 | 4.41 | 18.61 | 4.38 | 17.79 | 3.30 | 23.20 | 4.88 | 20.53 | 3.22 |
| **Slot** | *capacity K* | 1.27 | 1.44 | 1.37 | 0.35 | 0.88 | 0.51 | 1.30 | 0.35 | 1.88 | 0.23 | 1.88 | 0.24 | 1.81 | 0.41 | 1.87 | 0.26 |
|  | *sd* | 22.53 | 16.78 | 18.25 | 6.97 | 18.16 | 10.05 | 15.17 | 4.41 | 18.61 | 4.42 | 17.79 | 3.30 | 23.20 | 4.88 | 20.53 | 3.25 |
| **SR** | *capacity K* | 1.27 | 1.02 | 1.37 | 0.35 | 0.88 | 0.51 | 1.3 | 0.35 | 1.88 | 0.23 | 1.88 | 0.27 | 1.81 | 0.22 | 1.87 | 0.11 |
|  | *sd* | 22.74 | 11.69 | 16.33 | 6.78 | 18.76 | 8.44 | 13.68 | 4.23 | 13.79 | 3.34 | 13.11 | 2.50 | 17.30 | 4.39 | 15.09 | 2.61 |
| **EnsInt** | *g* | 0.4 | 0.31 | 0.31 | 0.18 | 0.56 | 0.27 | 0.35 | 0.18 | 0.06 | 0.11 | 0.06 | 0.10 | 0.09 | 0.11 | 0.06 | 0.06 |
|  | *sd* | 22.53 | 14.8 | 18.25 | 6.97 | 19.26 | 18.86 | 15.17 | 4.41 | 18.49 | 4.35 | 17.81 | 3.24 | 23.20 | 4.89 | 20.53 | 3.26 |
|  | *samples* | 7.82×10^12^ | 8.13×10^12^ | 6.39×10^12^ | 5.93×10^12^ | 5.96×10^12^ | 6.9×10^12^ | 6.86×10^12^ | 8.16×10^12^ | 3.07 | 1.84×10^12^ | 3.15 | 5.06×10^11^ | 2.12×10^12^ | 3.35×10^12^ | 9.15×10^11^ | 2.43×10^12^ |
| **Swap** | *g* | 0.36 | 0.28 | 0.31 | 0.18 | 0.53 | 0.26 | 0.31 | 0.18 | 0.06 | 0.09 | 0.05 | 0.08 | 0.09 | 0.11 | 0.06 | 0.06 |
|  | *B* | 2.02×10^-15^ | 0.05 | 2.97×10^-15^ | 0.01 | 2.89×10^-15^ | 5.33×10^-3^ | 2.31×10^-15^ | 5.06×10^-3^ | 5.05×10^-15^ | 0.02 | 9.64×10^-15^ | 0.02 | 3.59×10^-15^ | 1.28×10^-3^ | 5.84×10^-15^ | 3.44×10^-3^ |
|  | *sd* | 22.53 | 16.86 | 18.25 | 6.96 | 18.54 | 13.15 | 15.17 | 4.41 | 18.69 | 4.2 | 17.89 | 3.25 | 23.2 | 4.88 | 20.53 | 3.23 |
| **VPG** | *g* | 0.68 | 0.23 | 0.38 | 0.22 | 0.63 | 0.22 | 0.23 | 0.21 | 0.03 | 0.1 | 5.4×10^-3^ | 0.1 | 0.06 | 0.1 | 0.03 | 0.04 |
|  | *mode-*  *Precision* | 5×10^-4^ | 3.04×10^-3^ | 5×10^-4^ | 1.23×10^-4^ | 5×10^-4^ | 6.57×10^-4^ | 5.27×10^-4^ | 2.88×10^-4^ | 4.34×10^-3^ | 1.73×10^-3^ | 4.75×10^-3^ | 2.26×10^-3^ | 2.31×10^-3^ | 1.07×10^-3^ | 3.23×10^-3^ | 1.09×10^-3^ |
|  | *sdPrecision* | 6.69×10^-4^ | 10.42 | 5.01×10^-4^ | 7.72×10^-4^ | 5×10^-4^ | 6.11×10^-4^ | 5.01×10^-4^ | 2.84×10^-3^ | 4.38×10^-3^ | 3.39×10^-2^ | 4.9×10^-3^ | 8.86×10^-3^ | 1.92×10^-3^ | 2.02×10^-3^ | 2.82×10^-3^ | 2.04×10^-3^ |
| **VP** | *mode-*  *Precision* | 5×10^-4^ | 4.13×10^-12^ | 5×10^-4^ | 4.78×10^-5^ | 5×10^-4^ | 3.8×10^-12^ | 5×10^-4^ | 4.06×10^-5^ | 4.61×10^-3^ | 1.81×10^-3^ | 4.85×10^-3^ | 2.48×10^-3^ | 2.21×10^-3^ | 1.29×10^-3^ | 3.39×10^-3^ | 1.39×10^-3^ |
|  | *sd*Precision | 1.55×10^-3^ | 1.4×10^-4^ | 1.46×10^-3^ | 3.7×10^-4^ | 1.51×10^-3^ | 1.59×10^-4^ | 1.5×10^-3^ | 1.84×10^-3^ | 6.73×10^-3^ | 6.27×10^-3^ | 6.93×10^-3^ | 8.51×10^-3^ | 4.71×10^-3^ | 2.91×10^-3^ | 4.27×10^-3^ | 3.34×10^-3^ |

*(table continues)*

**Table S11** *(continued)*

|  |  | ***Exp3*** | | | | ***Exp4*** | | | | ***Exp5*** | | | |
| --- | --- | --- | --- | --- | --- | --- | --- | --- | --- | --- | --- | --- | --- |
|  |  | **Short SOA** | | **Long SOA** | | **Short SOA** | | **Long SOA** | | **Short SOA** | | **Long SOA** | |
| **Model** | **Parameter** | **Median** | **SD** | **Median** | **SD** | **Median** | **SD** | **Median** | **SD** | **Median** | **SD** | **Median** | **SD** |
| **StM** | *g* | 0.55 | 0.29 | 0.15 | 0.16 | 0.41 | 0.21 | 0.24 | 0.16 | 0.32 | 0.3 | 0.32 | 0.23 |
|  | *sd* | 17.65 | 13.67 | 18.45 | 6.18 | 16.13 | 7.59 | 15.77 | 4.67 | 18.14 | 13.58 | 16.93 | 6.39 |
| **Slot** | *capacity K* | 0.9 | 0.59 | 1.70 | 0.56 | 1.18 | 0.44 | 1.53 | 0.45 | 1.35 | 1.2 | 1.36 | 1.44 |
|  | *sd* | 17.65 | 13.67 | 18.45 | 6.17 | 16.13 | 7.59 | 15.77 | 4.94 | 18.14 | 13.45 | 17.08 | 6.37 |
| **SR** | *capacity K* | 0.9 | 0.62 | 1.70 | 0.83 | 1.18 | 0.46 | 1.53 | 0.61 | 1.35 | 0.78 | 1.36 | 0.47 |
|  | *sd* | 18.85 | 11.51 | 15.34 | 5.65 | 16.08 | 5.46 | 13.56 | 3.57 | 15.89 | 10.24 | 15.54 | 5.09 |
| **EnsInt** | *g* | 0.37 | 0.29 | 0.15 | 0.16 | 0.41 | 0.20 | 0.24 | 0.16 | 0.32 | 0.3 | 0.32 | 0.23 |
|  | *sd* | 21.38 | 15.34 | 18.45 | 6.18 | 16.00 | 7.18 | 15.77 | 4.11 | 18.02 | 13.9 | 16.93 | 6.39 |
|  | *samples* | 2.65×10^12^ | 1.09×10^13^ | 6.41×10^12^ | 8.49×10^12^ | 5.12×10^12^ | 6.55×10^12^ | 5.51×10^12^ | 5.92×10^12^ | 4.69×10^12^ | 7.77×10^12^ | 5.28×10^12^ | 1.05×10^13^ |
| **Swap** | *g* | 0.41 | 0.24 | 0.14 | 0.16 | 0.41 | 0.21 | 0.23 | 0.16 | 0.32 | 0.29 | 0.32 | 0.19 |
|  | *B* | 1.69×10^-12^ | 0.17 | 5.99×10^-15^ | 0.02 | 5.73×10^-15^ | 0.05 | 5.29×10^-15^ | 0.03 | 3.57×10^-15^ | 0.07 | 3.04×10^-15^ | 0.01 |
|  | *sd* | 18.32 | 11.66 | 18.45 | 6.15 | 16.13 | 6.73 | 16.08 | 4.52 | 18.51 | 13 | 16.93 | 7.23 |
| **VPG** | *g* | 0.66 | 0.28 | 0.14 | 0.19 | 0.44 | 0.20 | 0.21 | 0.16 | 0.61 | 0.3 | 0.36 | 0.23 |
|  | *mode-*  *Precision* | 5×10^-4^ | 0.36 | 5×10^-4^ | 2.58×10^-4^ | 5×10^-4^ | 2.1×10^-4^ | 5×10^-4^ | 1.66×10^-4^ | 5×10^-4^ | 0.82 | 5×10^-4^ | 1.61×10^-4^ |
|  | *sdPrecision* | 5×10^-4^ | 14.12 | 5×10^-4^ | 2.39×10^-4^ | 5.15×10^-4^ | 1.04×10^-3^ | 5.16×10^-4^ | 1.56×10^-3^ | 5.09×10^-4^ | 10.43 | 5.01×10^-4^ | 0.02 |
| **VP** | *mode-*  *Precision* | 5×10^-4^ | 2.58×10^-4^ | 5×10^-4^ | 2.02×10^-4^ | 5×10^-4^ | 8.05×10^-12^ | 5×10^-4^ | 6.64×10^-5^ | 5×10^-4^ | 5.85×10^-5^ | 5×10^-4^ | 1.02×10^-4^ |
|  | *mode*-  *sdPrecision* | 1.54×10^-3^ | 4.22×10^-4^ | 1.08×10^-3^ | 1.24×10^-3^ | 1.5×10^-3^ | 1.9×10^-4^ | 1.5×10^-3^ | 1.04×10^-3^ | 1.45×10^-3^ | 5.96×10^-4^ | 1.48×10^-3^ | 5.15×10^-4^ |

**Table S12**

*Maximum-likelihood estimates of the parameters in all models in the T1 incorrect condition.*

|  |  | ***Exp1A*** | | | | ***Exp1B*** | | | | ***Exp2A*** | | | | ***Exp2B*** | | | |
| --- | --- | --- | --- | --- | --- | --- | --- | --- | --- | --- | --- | --- | --- | --- | --- | --- | --- |
|  |  | **Short SOA** | | **Long SOA** | | **Short SOA** | | **Long SOA** | | **Short SOA** | | **Long SOA** | | **Short SOA** | | **Long SOA** | |
| **Model** | **Parameter** | **Median** | **SD** | **Median** | **SD** | **Median** | **SD** | **Median** | **SD** | **Median** | **SD** | **Median** | **SD** | **Median** | **SD** | **Median** | **SD** |
| **StM** | *g* | 0.89 | 0.22 | 0.71 | 0.26 | 0.88 | 0.25 | 0.54 | 0.24 | 0.29 | 0.26 | 0.21 | 0.26 | 0.46 | 0.25 | 0.30 | 0.25 |
|  | *sd* | 14.44 | 11.99 | 13.76 | 10.74 | 15.91 | 8.06 | 13.67 | 8.09 | 15.98 | 22.54 | 16.35 | 26.11 | 20.66 | 15.65 | 20.22 | 12.22 |
| **Slot** | *capacity K* | 0.15 | 0.26 | 0.56 | 0.47 | 0.23 | 0.5 | 0.93 | 0.51 | 1.42 | 0.90 | 1.58 | 1.49 | 1.08 | 0.76 | 1.39 | 0.81 |
|  | *sd* | 11.44 | 11.72 | 13.12 | 8.01 | 16.18 | 34.64 | 13.67 | 8.14 | 15.08 | 8.20 | 16.35 | 26.11 | 20.66 | 15.65 | 17.61 | 12.91 |
| **SR** | *capacity K* | 0.21 | 0.43 | 0.56 | 0.47 | 0.24 | 0.53 | 0.93 | 0.49 | 1.42 | 0.92 | 1.58 | 0.80 | 1.20 | 0.62 | 1.39 | 0.62 |
|  | *sd* | 22.33 | 35.28 | 15.15 | 10.53 | 20.74 | 57.87 | 14.39 | 7.45 | 13.95 | 8.02 | 13.41 | 20.44 | 21.65 | 17.79 | 17.86 | 10.19 |
| **EnsInt** | *g* | 0.8 | 0.41 | 0.64 | 0.3 | 0.69 | 0.39 | 0.53 | 0.24 | 0.29 | 0.28 | 0.28 | 0.30 | 0.32 | 0.24 | 0.29 | 0.29 |
|  | *sd* | 22.25 | 22.79 | 15.72 | 14.13 | 16.27 | 28.57 | 14.16 | 11.1 | 15.13 | 12.29 | 13.43 | 9.30 | 22.02 | 25.40 | 17.61 | 14.80 |
|  | *samples* | 0.47 | 1.29×10^12^ | 10.42 | 1.35×10^12^ | 1.65 | 1.26×10^13^ | 6.78×10^11^ | 3.21×10^12^ | 0.51 | 1.33×10^12^ | 0.56 | 3.52×10^12^ | 0.82 | 1.42×10^12^ | 1.04 | 2.81×10^12^ |
| **Swap** | *g* | 0.72 | 0.34 | 0.47 | 0.27 | 0.69 | 0.33 | 0.48 | 0.23 | 0.14 | 0.22 | 1.09×10^-10^ | 0.19 | 0.13 | 0.17 | 0.12 | 0.22 |
|  | *B* | 0.05 | 0.28 | 0.04 | 0.23 | 0.02 | 0.17 | 0.02 | 0.06 | 2.25×10^-8^ | 0.13 | 0.04 | 0.15 | 0.16 | 0.18 | 0.09 | 0.10 |
|  | *sd* | 17.36 | 18.91 | 16.31 | 14.21 | 16.27 | 66.22 | 14.49 | 7.14 | 15.92 | 22.18 | 18.01 | 24.58 | 24.70 | 15.22 | 20.09 | 11.45 |
| **VPG** | *g* | 0.94 | 0.12 | 0.75 | 0.26 | 0.87 | 0.2 | 0.55 | 0.25 | 0.28 | 0.29 | 0.26 | 0.30 | 0.38 | 0.25 | 0.29 | 0.25 |
|  | *mode-*  *Precision* | 6.01×10^-4^ | 1.28 | 7.39×10^-4^ | 0.02 | 5.55×10^-4^ | 2.55×10^-3^ | 5×10^-4^ | 1.63×10^-3^ | 4.61×10^-3^ | 0.90 | 3.65×10^-3^ | 0.01 | 1.47×10^-3^ | 4.71×10^-3^ | 2.28×10^-3^ | 0.08 |
|  | *sdPrecision* | 5.02×10^-4^ | 14.38 | 5×10^-4^ | 10.47 | 5×10^-4^ | 3.9×10^-3^ | 6.07×10^-4^ | 8.24×10^-3^ | 1×10^-3^ | 19.61 | 1×10^-3^ | 0.01 | 1×10^-3^ | 0.03 | 1×10^-3^ | 26.69 |
| **VP** | *mode-*  *Precision* | 5×10^-4^ | 5.35×10^-12^ | 5×10^-4^ | 8.88×10^-11^ | 5×10^-4^ | 6.14×10^-12^ | 5×10^-4^ | 1.55×10^-11^ | 1×10^-3^ | 0.04 | 1×10^-3^ | 3.34×10^-3^ | 1×10^-3^ | 2.76×10^-3^ | 1×10^-3^ | 3.48×10^-3^ |
|  | *sd*Precision | 1.64×10^-3^ | 1.04×10^-4^ | 1.66×10^-3^ | 3.15×10^-4^ | 1.64×10^-3^ | 2.11×10^-4^ | 1.65×10^-3^ | 4.53×10^-4^ | 4.56×10^-3^ | 0.32 | 4.49×10^-3^ | 0.01 | 3.71×10^-3^ | 0.01 | 3.52×10^-3^ | 0.03 |

*(table continues)*

**Table S12** *(continued)*

|  |  | ***Exp3*** | | | | ***Exp4*** | | | | ***Exp5*** | | | |
| --- | --- | --- | --- | --- | --- | --- | --- | --- | --- | --- | --- | --- | --- |
|  |  | **Short SOA** | | **Long SOA** | | **Short SOA** | | **Long SOA** | | **Short SOA** | | **Long SOA** | |
| **Model** | **Parameter** | **Median** | **SD** | **Median** | **SD** | **Median** | **SD** | **Median** | **SD** | **Median** | **SD** | **Median** | **SD** |
| **StM** | *g* | 0.72 | 0.33 | 0.43 | 0.32 | 0.54 | 0.25 | 0.43 | 0.19 | 0.61 | 0.31 | 0.59 | 0.3 |
|  | *sd* | 15.35 | 16.54 | 17.29 | 10.80 | 14.55 | 10.36 | 15.49 | 7.21 | 18.95 | 13.59 | 15.24 | 17.52 |
| **Slot** | *capacity K* | 0.56 | 0.73 | 1.13 | 0.82 | 0.92 | 0.53 | 1.14 | 0.41 | 0.72 | 0.64 | 0.77 | 1.64 |
|  | *sd* | 13.76 | 16.75 | 17.29 | 12.45 | 14.55 | 10.36 | 15.49 | 7.21 | 18.16 | 26.34 | 13.41 | 17.53 |
| **SR** | *capacity K* | 0.56 | 0.82 | 1.13 | 0.95 | 0.92 | 0.79 | 1.14 | 0.41 | 0.97 | 0.75 | 0.75 | 1.12 |
|  | *sd* | 20.53 | 12.36 | 15.54 | 9.26 | 16.28 | 7.75 | 14.72 | 5.50 | 23.33 | 42.1 | 15.22 | 16.46 |
| **EnsInt** | *g* | 0.68 | 0.33 | 0.53 | 0.32 | 0.51 | 0.25 | 0.43 | 0.19 | 0.44 | 0.33 | 0.59 | 0.31 |
|  | *sd* | 13.76 | 18.90 | 13.92 | 8.12 | 14.22 | 10.92 | 15.51 | 7.19 | 22.21 | 45.71 | 14.89 | 13.3 |
|  | *samples* | 2.19 | 8.51×10^12^ | 2.12×10^11^ | 5.21×10^12^ | 3.85×10^11^ | 5.11×10^12^ | 1.65×10^12^ | 5×10^12^ | 1.23×10^12^ | 6.22×10^12^ | 4.27×10^11^ | 5.57×10^12^ |
| **Swap** | *g* | 0.35 | 0.31 | 0.3 | 0.33 | 0.4 | 0.24 | 0.33 | 0.19 | 0.44 | 0.34 | 0.51 | 0.31 |
|  | *B* | 0.08 | 0.23 | 1.14×10^-10^ | 0.20 | 0.05 | 0.08 | 0.04 | 0.09 | 1.75×10^-3^ | 0.25 | 3.41×10^-14^ | 0.29 |
|  | *sd* | 25.61 | 46.16 | 19.65 | 13.54 | 14.61 | 9.95 | 15.36 | 6.91 | 20.96 | 59.92 | 16.00 | 14.67 |
| **VPG** | *g* | 0.84 | 0.24 | 0.49 | 0.34 | 0.54 | 0.19 | 0.40 | 0.18 | 0.75 | 0.3 | 0.61 | 0.29 |
|  | *mode-*  *Precision* | 6.35×10^-4^ | 0.01 | 5.67×10^-4^ | 0.03 | 5×10^-4^ | 1.22×10^-3^ | 5.04×10^-4^ | 3.52×10^-4^ | 5.04×10^-4^ | 5.82×10^-4^ | 5.15×10^-4^ | 4.05×10^-3^ |
|  | *sdPrecision* | 5.75×10^-4^ | 10.2 | 5×10^-4^ | 4.75×10^-3^ | 5.25×10^-4^ | 0.91 | 5.11×10^-4^ | 0.01 | 1.07×10^-3^ | 0.02 | 5×10^-4^ | 0.02 |
| **VP** | *mode-*  *Precision* | 5×10^-4^ | 8.89×10^-12^ | 5×10^-4^ | 7.49×10^-4^ | 5×10^-4^ | 9.77×10^-12^ | 5×10^-4^ | 1.23×10^-11^ | 5×10^-4^ | 1.99×10^-11^ | 5×10^-4^ | 1.03×10^-4^ |
|  | *mode*-  *sdPrecision* | 1.62×10^-3^ | 2.78×10^-4^ | 1.47×10^-3^ | 2.35×10^-3^ | 1.58×10^-3^ | 2.93×10^-4^ | 1.52×10^-3^ | 4.32×10^-4^ | 1.59×10^-3^ | 6.02×10^-4^ | 1.58×10^-3^ | 0.01 |

**4. Wang et al.**

**Table S13**

*Model comparison results for T2 performance in the T1 correct condition (T2|T1).*

|  |  | **CC** | | | | **CO** | | | | **OC** | | | | **OO** | | | |
| --- | --- | --- | --- | --- | --- | --- | --- | --- | --- | --- | --- | --- | --- | --- | --- | --- | --- |
| **Lag** | **Model** | **MeanAIC** | **Mean ΔAIC** | **Mean**  **Rank** | **Mean**  **Log-Likelihood** | **MeanAIC** | **Mean**  **ΔAIC** | **Mean**  **Rank** | **Mean**  **Log-Likelihood** | **MeanAIC** | **Mean ΔAIC** | **Mean Rank** | **Mean**  **Log-Likelihood** | **MeanAIC** | **Mean ΔAIC** | **Mean Rank** | **Mean**  **Log-Likelihood** |
| **1** | StM | 277.06 | 1.25 | 3.16 | -136.53 | 338.47 | 0.47 | 2.36 | -167.23 | 233.70 | 1.01 | 3.25 | -114.85 | 124.84 | 1.29 | 2.82 | -60.42 |
|  | Slot | 277.06 | 1.25 | 2.88 | -136.53 | 338.48 | 0.48 | 2.41 | -167.24 | 233.70 | 1.01 | 2.43 | -114.85 | 124.73 | 1.19 | 2.48 | -60.37 |
|  | SR | 277.98 | 2.17 | 3.39 | -136.99 | 338.50 | 0.50 | 2.41 | -167.25 | 234.13 | 1.44 | 2.75 | -115.06 | 124.84 | 1.29 | 2.41 | -60.42 |
|  | EnsInt | 278.17 | 2.36 | 4.84 | -136.09 | 340.17 | 2.17 | 5.20 | -167.09 | 234.57 | 1.88 | 4.79 | -114.29 | 125.37 | 1.83 | 4.36 | -59.69 |
|  | Swap | 277.24 | 1.43 | 4.30 | -135.62 | 340.47 | 2.47 | 5.55 | -167.23 | 235.63 | 2.94 | 6.32 | -114.82 | 125.35 | 1.81 | 4.61 | -59.68 |
|  | VPG | 278.74 | 2.93 | 5.82 | -136.37 | 340.63 | 2.63 | 5.75 | -167.32 | 235.24 | 2.55 | 5.82 | -114.62 | 127.54 | 3.99 | 6.39 | -60.77 |
|  | VP | 277.86 | 2.05 | 3.61 | -136.93 | 342.66 | 4.66 | 4.32 | -169.33 | 233.16 | 0.47 | 2.64 | -114.58 | 127.97 | 4.42 | 4.93 | -61.98 |
| **3** | StM | 371.72 | 1.64 | 3.00 | -183.86 | 371.13 | 0.44 | 2.41 | -183.57 | 219.47 | 1.31 | 3.07 | -107.73 | 309.69 | 3.60 | 2.73 | -152.85 |
|  | Slot | 371.72 | 1.64 | 2.79 | -183.86 | 371.03 | 0.33 | 2.21 | -183.51 | 219.47 | 1.31 | 2.61 | -107.73 | 309.65 | 3.56 | 2.48 | -152.83 |
|  | SR | 375.11 | 5.03 | 4.36 | -185.56 | 371.05 | 0.36 | 2.23 | -183.53 | 220.51 | 2.35 | 3.25 | -108.25 | 309.56 | 3.46 | 2.29 | -152.78 |
|  | EnsInt | 373.23 | 3.15 | 5.30 | -183.62 | 372.61 | 1.92 | 4.55 | -183.31 | 220.83 | 2.67 | 5.00 | -107.41 | 309.87 | 3.77 | 3.86 | -151.93 |
|  | Swap | 373.03 | 2.95 | 5.38 | -183.52 | 372.41 | 1.71 | 4.13 | -183.20 | 221.32 | 3.17 | 5.93 | -107.66 | 308.52 | 2.43 | 4.07 | -151.26 |
|  | VPG | 372.37 | 2.29 | 4.75 | -183.19 | 373.74 | 3.05 | 5.68 | -183.87 | 220.52 | 2.36 | 5.18 | -107.26 | 312.17 | 6.08 | 5.68 | -153.09 |
|  | VP | 371.08 | 1.00 | 2.43 | -183.54 | 397.39 | 26.70 | 6.79 | -196.70 | 218.74 | 0.59 | 2.96 | -107.37 | 336.39 | 30.30 | 6.89 | -166.20 |
| **8** | StM | 378.88 | 1.90 | 3.07 | -187.44 | 347.09 | 0.59 | 2.30 | -171.55 | 220.87 | 1.29 | 3.23 | -108.43 | 331.65 | 1.24 | 2.23 | -163.82 |
|  | Slot | 378.89 | 1.91 | 2.86 | -187.44 | 347.09 | 0.59 | 2.30 | -171.55 | 221.25 | 1.67 | 2.80 | -108.63 | 331.65 | 1.24 | 2.23 | -163.82 |
|  | SR | 380.89 | 3.92 | 3.68 | -188.45 | 347.09 | 0.59 | 2.25 | -171.55 | 221.10 | 1.51 | 3.04 | -108.55 | 331.65 | 1.24 | 2.29 | -163.82 |
|  | EnsInt | 380.41 | 3.43 | 5.25 | -187.20 | 348.38 | 1.87 | 4.39 | -171.19 | 221.87 | 2.29 | 4.71 | -107.93 | 333.17 | 2.76 | 4.46 | -163.58 |
|  | Swap | 380.28 | 3.30 | 5.50 | -187.14 | 348.69 | 2.18 | 4.75 | -171.34 | 222.76 | 3.18 | 6.50 | -108.38 | 332.16 | 1.76 | 4.46 | -163.08 |
|  | VPG | 379.39 | 2.42 | 4.96 | -186.70 | 349.22 | 2.71 | 5.61 | -171.61 | 222.12 | 2.54 | 5.25 | -108.06 | 334.13 | 3.72 | 6.11 | -164.06 |
|  | VP | 377.73 | 0.75 | 2.68 | -186.86 | 363.37 | 16.86 | 6.39 | -179.68 | 220.38 | 0.79 | 2.46 | -108.19 | 343.72 | 13.31 | 6.21 | -169.86 |

*Note.* CC = color-color target pair; CO = color-orientation target pair; OC = orientation-color target pair; OO = orientation-orientation target pair. See *2.1. Model details* of the main text for the model abbreviations.

**Table S14**

*Model comparison results for T2 performance in the T1 incorrect condition.*

|  | **Lag 1** | | | | **Lag 3** | | | | **Lag 8** | | | |
| --- | --- | --- | --- | --- | --- | --- | --- | --- | --- | --- | --- | --- |
| **Model** | **MeanAIC** | **Mean ΔAIC** | **Mean**  **Rank** | **Mean**  **Log-Likelihood** | **MeanAIC** | **Mean**  **ΔAIC** | **Mean**  **Rank** | **Mean**  **Log-Likelihood** | **MeanAIC** | **Mean ΔAIC** | **Mean Rank** | **Mean**  **Log-Likelihood** |
| StM | 614.14 | 12.62 | 3.61 | -305.068 | 336.86 | 0.95 | 2.46 | -166.429 | 277.62 | 0.63 | 2.59 | -136.808 |
| Slot | 613.96 | 12.45 | 3.34 | -304.982 | 336.88 | 0.97 | 2.61 | -166.44 | 277.58 | 0.60 | 2.52 | -136.791 |
| SR | 613.87 | 12.35 | 3.30 | -304.934 | 336.88 | 0.97 | 2.61 | -166.44 | 277.62 | 0.63 | 2.54 | -136.808 |
| EnsInt | 613.65 | 12.14 | 4.04 | -303.825 | 338.27 | 2.36 | 4.38 | -166.135 | 278.62 | 1.63 | 4.34 | -136.309 |
| Swap | 601.55 | 0.04 | 1.11 | -297.775 | 337.24 | 1.33 | 3.77 | -165.618 | 279.29 | 2.31 | 5.16 | -136.645 |
| VPG | 616.12 | 14.61 | 5.71 | -305.061 | 338.91 | 3.01 | 5.54 | -166.457 | 279.04 | 2.05 | 5.00 | -136.519 |
| VP | 652.02 | 50.51 | 6.89 | -324.01 | 352.78 | 16.88 | 6.64 | -174.392 | 286.00 | 9.01 | 5.86 | -140.998 |

*Note.* For the T1 incorrect condition, analyses were conducted for each specific lag, across all target pairs. See the main text for a more detailed discussion.

**Table S15**

*Maximum-likelihood estimates of the parameters in all models in the T1 correct condition.*

|  |  | **CC** | | | | | | **CO** | | | | | |
| --- | --- | --- | --- | --- | --- | --- | --- | --- | --- | --- | --- | --- | --- |
|  |  | **Lag 1** | | **Lag 3** | | **Lag 8** | | **Lag 1** | | **Lag 3** | | **Lag 8** | |
| **Model** | **Parameter** | **Median** | **SD** | **Median** | **SD** | **Median** | **SD** | **Median** | **SD** | **Median** | **SD** | **Median** | **SD** |
| **StM** | *g* | 0.07 | 0.11 | 0.04 | 0.07 | 0.06 | 0.08 | 0.06 | 0.14 | 0.33 | 0.33 | 0.29 | 0.30 |
|  | *sd* | 9.13 | 2.83 | 8.84 | 1.71 | 8.95 | 1.73 | 14.61 | 9.88 | 17.71 | 17.29 | 15.38 | 6.03 |
| **Slot** | *capacity K* | 1.86 | 2.36 | 1.91 | 1.45 | 1.88 | 0.92 | 1.87 | 1.17 | 1.32 | 1.16 | 1.41 | 0.60 |
|  | *sd* | 9.13 | 2.83 | 8.84 | 1.71 | 8.95 | 1.77 | 14.88 | 9.87 | 15.67 | 16.82 | 14.70 | 8.63 |
| **SR** | *capacity K* | 2.20 | 1.01 | 2.61 | 1.02 | 2.59 | 0.83 | 1.90 | 0.86 | 1.32 | 0.90 | 1.44 | 0.78 |
|  | *sd* | 6.84 | 1.98 | 7.62 | 1.20 | 6.99 | 1.13 | 11.16 | 6.81 | 18.86 | 11.81 | 13.13 | 31.61 |
| **EnsInt** | *g* | 0.07 | 0.11 | 0.04 | 0.07 | 0.06 | 0.08 | 0.09 | 0.24 | 0.29 | 0.34 | 0.30 | 0.29 |
|  | *sd* | 8.89 | 2.82 | 8.73 | 1.74 | 8.75 | 1.69 | 13.82 | 6.25 | 19.74 | 19.33 | 13.13 | 7.78 |
|  | *samples* | 4.27 | 2.24×10^12^ | 5.78×10^11^ | 3.18×10^12^ | 5.06×10^10^ | 2×10^12^ | 21.42 | 2,18×10^12^ | 2.78×10^11^ | 4.54×10^12^ | 9.25 | 2.57×10^12^ |
| **Swap** | *g* | 6.37×10^-8^ | 0.09 | 0.04 | 0.07 | 0.02 | 0.08 | 0.06 | 0.12 | 0.23 | 0.29 | 0.27 | 0.29 |
|  | *B* | 0.02 | 0.05 | 1.52×10^-11^ | 0.02 | 4.36×10^-13^ | 0.02 | 5.42×10^-7^ | 0.07 | 4.8×10^-3^ | 0.13 | 7.36×10^-14^ | 0.04 |
|  | *sd* | 9.20 | 2.37 | 8.82 | 1.57 | 8.95 | 2.06 | 14.74 | 8.46 | 20.87 | 16.49 | 15.22 | 15.37 |
| **VPG** | *g* | 0.03 | 0.11 | 5.85×10^-6^ | 0.07 | 8.16×10^-4^ | 0.06 | 0.07 | 0.22 | 0.66 | 0.26 | 0.29 | 0.3 |
|  | *mode-*  *Precision* | 1.73×10^-3^ | 1.13×10^-3^ | 1.97×10^-3^ | 1.54×10^-3^ | 1.81×10^-3^ | 3.82×10^-3^ | 6.75×10^-4^ | 5.19×10^-4^ | 5.22×10^-4^ | 0.17 | 6.24×10^-4^ | 7.67×10^-4^ |
|  | *sdPrecision* | 9.38×10^-4^ | 0.01 | 1.1×10^-3^ | 0.03 | 1.8×10^-3^ | 0.03 | 5.01×10^-4^ | 6.31×10^-3^ | 5×10^-4^ | 13.11 | 5×10^-4^ | 8.06×10^-3^ |
| **VP** | *mode-*  *Precision* | 1.64×10^-3^ | 9.67×10^-4^ | 2.25×10^-3^ | 1.58×10^-3^ | 1.97×10^-3^ | 2.04×10^-3^ | 5.44×10^-4^ | 5.21×10^-4^ | 5×10^-4^ | 1.73×10^-4^ | 5×10^-4^ | 4.47×10^-4^ |
|  | *sdPrecision* | 2.63×10^-3^ | 0.01 | 3.47×10^-3^ | 0.03 | 3.88×10^-3^ | 7.02×10^-3^ | 1.12×10^-3^ | 5.99×10^-3^ | 1.52×10^-3^ | 1.15×10^-3^ | 1.68×10^-3^ | 5.56×10^-3^ |

*(table continues)*

**Table S15** *(continued)*

|  |  | **OC** | | | | | | **OO** | | | | | |
| --- | --- | --- | --- | --- | --- | --- | --- | --- | --- | --- | --- | --- | --- |
|  |  | **Lag 1** | | **Lag 3** | | **Lag 8** | | **Lag 1** | | **Lag 3** | | **Lag 8** | |
| **Model** | **Parameter** | **Median** | **SD** | **Median** | **SD** | **Median** | **SD** | **Median** | **SD** | **Median** | **SD** | **Median** | **SD** |
| **StM** | *g* | 2.05×10^-14^ | 0.04 | 1.12×10^-13^ | 0.08 | 3.24×10^-14^ | 0.06 | 6.28×10^-14^ | 0.26 | 0.51 | 0.38 | 0.28 | 0.30 |
|  | *sd* | 7.75 | 2.62 | 7.73 | 2.60 | 7.43 | 2.43 | 15.54 | 10.26 | 15.91 | 41.36 | 18.56 | 10.40 |
| **Slot** | *capacity K* | 2.43 | 2.32 | 2.05 | 2.01 | 2.35 | 2.35 | 2.12 | 1.51 | 0.99 | 1.17 | 1.43 | 1.67 |
|  | *sd* | 7.75 | 2.62 | 7.73 | 2.60 | 7.43 | 2.67 | 15.37 | 10.61 | 13.84 | 25.95 | 18.56 | 12.42 |
| **SR** | *capacity K* | 2.76 | 0.85 | 2.74 | 1.00 | 2.74 | 0.90 | 2.05 | 1.22 | 1.12 | 1.01 | 1.43 | 0.62 |
|  | *sd* | 5.61 | 1.85 | 6.14 | 1.84 | 5.93 | 1.86 | 13.26 | 6.89 | 17.77 | 60.31 | 15.68 | 50.19 |
| **EnsInt** | *g* | 1.13×10^-14^ | 0.06 | 0.02 | 0.08 | 6.41×10^-15^ | 0.06 | 1.29×10^-4^ | 0.22 | 0.46 | 0.36 | 0.28 | 0.27 |
|  | *sd* | 7.75 | 2.72 | 7.54 | 2.54 | 7.35 | 2.54 | 15.93 | 9.89 | 19.87 | 44.40 | 18.32 | 7.95 |
|  | *samples* | 1.36 | 1.8×10^12^ | 2.09 | 2.25×10^12^ | 14.07 | 2.78×10^12^ | 8.96 | 8.72×10^12^ | 0.47 | 5.85×10^12^ | 6.52×10^12^ | 8.99×10^12^ |
| **Swap** | *g* | 2.37×10^-9^ | 0.03 | 1.45×10^-5^ | 0.07 | 3.04×10^-8^ | 0.06 | 1.93×10^-9^ | 0.21 | 0.29 | 0.27 | 0.22 | 0.22 |
|  | *B* | 9.58×10^-14^ | 0.03 | 2.65×10^-14^ | 0.02 | 3.14×10^-14^ | 0.01 | 8.95×10^-6^ | 0.23 | 7.27×10^-6^ | 0.32 | 2.26×10^-14^ | 0.25 |
|  | *sd* | 7.75 | 2.42 | 7.73 | 2.59 | 7.46 | 2.44 | 14.12 | 8.90 | 14.34 | 44.83 | 16.16 | 13.24 |
| **VPG** | *g* | 1.15×10^-8^ | 0.03 | 2.38×10^-4^ | 0.05 | 2.38×10^-6^ | 0.04 | 0.03 | 0.28 | 0.69 | 0.27 | 0.41 | 0.31 |
|  | *mode-*  *Precision* | 3.05×10^-3^ | 5.49×10^-3^ | 2.82×10^-3^ | 7.39×10^-3^ | 2.97×10^-3^ | 2.86 | 6.67×10^-4^ | 0.03 | 6.79×10^-4^ | 5.78×10^-3^ | 5.08×10^-4^ | 1.68×10^-4^ |
|  | *sdPrecision* | 9.36×10^-4^ | 3.38×10^-3^ | 1.08×10^-3^ | 0.03 | 1.73×10^-3^ | 9.45 | 5.19×10^-4^ | 0.1 | 5×10^-4^ | 9.45 | 5×10^-4^ | 6.93×10^-4^ |
| **VP** | *mode-*  *Precision* | 2.54×10^-3^ | 3.27×10^-3^ | 2.81×10^-3^ | 3.41×10^-3^ | 2.96×10^-3^ | 1.41 | 5×10^-4^ | 2.13×10^-4^ | 5×10^-4^ | 2.01×10^-5^ | 5×10^-4^ | 1.02×10^-4^ |
|  | *sdPrecision* | 9.15×10^-4^ | 0.02 | 2.92×10^-3^ | 3.04×10^-2^ | 1.74×10^-3^ | 9.45 | 9.6×10^-4^ | 5.13×10^-3^ | 1.6×10^-3^ | 4.77×10^-4^ | 1.37×10^-3^ | 3.59×10^-4^ |

**Table S16**

*Maximum-likelihood estimates of the parameters in all models in the T1 incorrect condition.*

|  |  | **Lag 1** | | **Lag 3** | | **Lag 8** | |
| --- | --- | --- | --- | --- | --- | --- | --- |
| **Model** | **Parameter** | **Median** | **SD** | **Median** | **SD** | **Median** | **SD** |
| **StM** | *g* | 0.54 | 0.29 | 0.60 | 0.26 | 0.33 | 0.20 |
|  | *sd* | 15.56 | 9.91 | 9.64 | 7.76 | 10.88 | 5.11 |
| **Slot** | *capacity K* | 0.93 | 0.56 | 0.80 | 1.37 | 1.24 | 0.48 |
|  | *sd* | 14.12 | 36.64 | 9.64 | 7.76 | 10.77 | 5.47 |
| **SR** | *capacity K* | 0.93 | 0.56 | 0.80 | 0.78 | 1.33 | 0.51 |
|  | *sd* | 14.12 | 36.64 | 11.75 | 10.20 | 9.58 | 3.88 |
| **EnsInt** | *g* | 0.33 | 0.28 | 0.55 | 0.28 | 0.33 | 0.20 |
|  | *sd* | 18.25 | 14.46 | 10.03 | 10.45 | 10.08 | 5.64 |
|  | *samples* | 0.97 | 2.48×10^12^ | 8.29 | 1.83×10^12^ | 3.51×10^11^ | 5.82×10^12^ |
| **Swap** | *g* | 0.22 | 0.18 | 0.38 | 0.23 | 0.26 | 0.19 |
|  | *B* | 0.26 | 0.13 | 0.04 | 0.14 | 4.87×10^-14^ | 0.07 |
|  | *sd* | 15.30 | 6.33 | 10.69 | 10.48 | 11.97 | 5.29 |
| **VPG** | *g* | 0.57 | 0.24 | 0.55 | 0.24 | 0.35 | 0.19 |
|  | *mode-*  *Precision* | 5.09×10^-4^ | 2.14 | 1.13×10^-3^ | 2.42×10^-3^ | 1.15×10^-3^ | 5.25×10^-3^ |
|  | *sdPrecision* | 5.68×10^-4^ | 9.44 | 5×10^-4^ | 9.45 | 5×10^-4^ | 4.23 |
| **VP** | *mode-*  *Precision* | 5×10^-4^ | 2.15×10^-11^ | 5×10^-4^ | 2.3×10^-9^ | 5×10^-4^ | 2.69×10^-4^ |
|  | *sdPrecision* | 1.64×10^-3^ | 8.26×10^-4^ | 1.97×10^-3^ | 0.01 | 2.37×10^-3^ | 9.22×10^-3^ |

**B. Additional analysis for the Ensemble Integration (EnsInt) and Swap models, using the data from Tang et al. (2022).**

We performed an extended analysis of the EnsInt and Swap models using the data from Tang et al. (2022), aiming to examine the potential interactions between the target (e.g., T2) and the subsequent mask. Specifically, we introduced three versions for these two models: 1) With-T1. This is the original model discussed in the main text, which accounts for T1 as the sole distractor in the analysis. 2) With-T2mask. This model uniquely considers the subsequent mask as the distractor. 3) With-Both. This model incorporates both T1 and the subsequent mask as the distractors of T2. The methodologies for the model fitting and comparison are consistent with those described in *2. Method* of the main text*.*

In the analysis of the EnsInt model (Figure S1), we observed different patterns between Experiment 1 and Experiment 2. For Experiment 1, the With-T1 version exhibited the best fitness to the data at shorter lags (Lag 1, 2 and 3), compared to longer lags (Lag 5 and 7). In contrast, the With-T2mask version’s performance improved with increasing lag, as indicated at both the subject and group levels. This trend may be attributed to the growing temporal distance between T1 and T2; at longer lags, T2 is more influenced by the subsequent mask than by T1. For Experiment 2, the With-T2mask version showed better fitness both at Lag 3 and Lag 7. Generally, the differences of AIC values are relatively minor (within the range from 0 to 10).

In the analysis of the Swap model (Figure S2), the difference among these three variants were slightly larger, compared to those observed in the EnsInt model. For Experiment 1, it was clearly observed that the With-T1 version consistently showed a superior performance across different lags, although the With-T2mask’s fitness improved as lag increased. This suggests that as the time interval between two targets increases, the likelihood of swaps between T2 and its subsequent mask increases. However, despite the observed increase in fitness for the With-T2mask version at longer lags, we still believe that T1 is likely to be a predominant distractor to T2, with regard to potential swap errors. For Experiment 2, the differences among these versions were small at Lag 3. At Lag 7, the With-T2mask showed a slightly better performance than others, which was consistent with the trend observed in the EnsInt model analysis.

Considering the relatively small differences among the variants of both the EnsInt and Swap models, along with the good performance of the With-T1 version of the model, we chose to present this version in the main text for simplicity. It is important to note, however, that our choice did not rule out the potential influence of the subsequent mask’s interaction. Unfortunately, due to the design of other datasets, where no identical item types followed T2, we are unable to conduct a similar extended analysis in those cases. Our focus on the With-T1 version is thus a reflection of both the data available, and our aim to maintain a streamlined presentation.

**Figure S1**

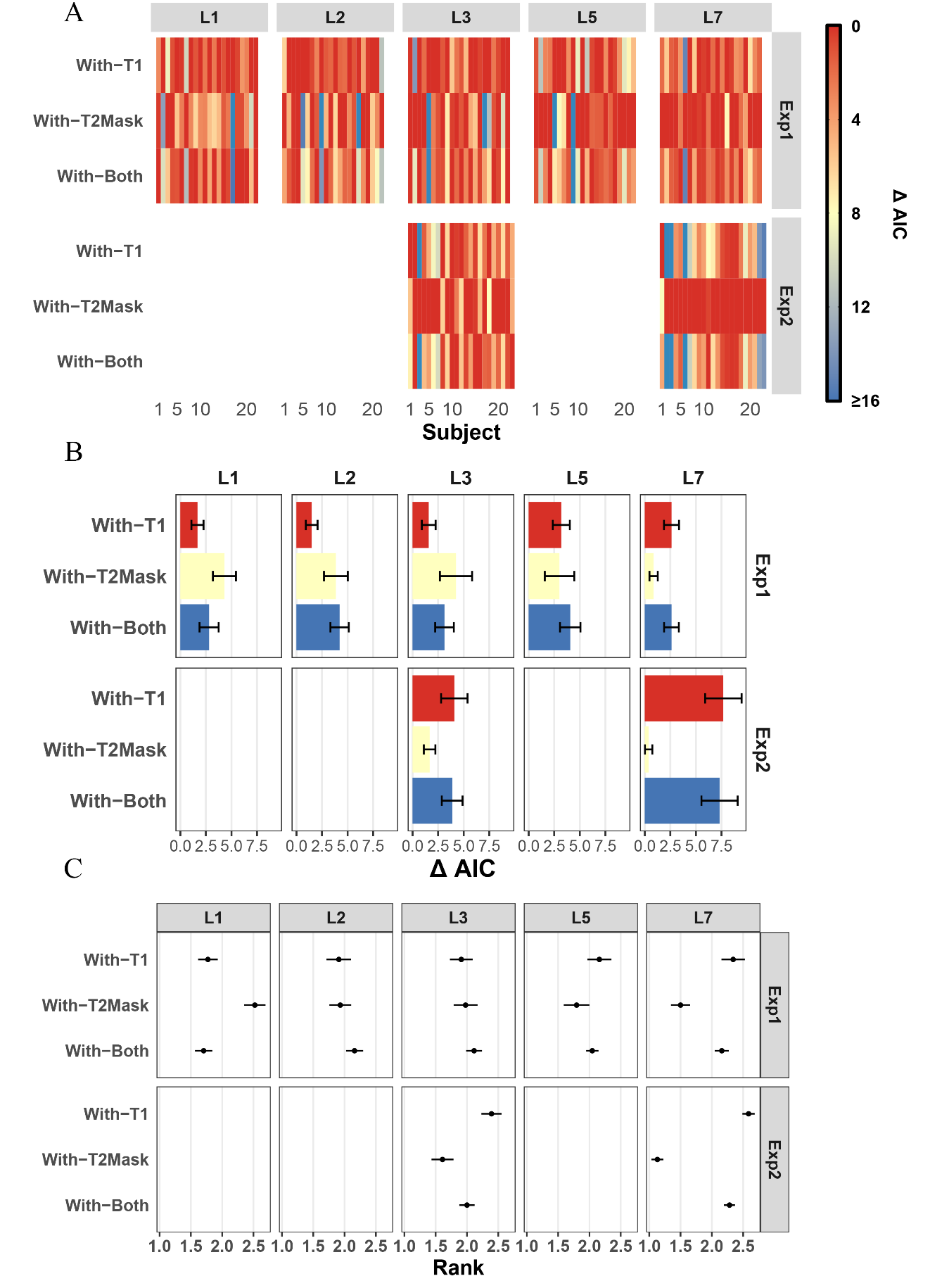
*Model comparison results for the EnsInt model and its variants.*

*Note.* With-T1 = only T1 is considered as the distractor. With-T2Mask = only the subsequent mask of T2 is considered as the distractor. With-Both = both T1 and the subsequent mask of T2 are considered as the distractors. L1 = Lag 1; L2 = Lag 2; L3 = Lag 3; L5 = Lag 5; L7 = Lag7. Exp = Experiment. (**A**) Model comparison results at the subject level, using the data from Tang et al. (2020) (**B**) Relative AIC values compared to that of the best-performing model, averaged across all subjects. (**C**) Average model ranking.

**Figure S2**

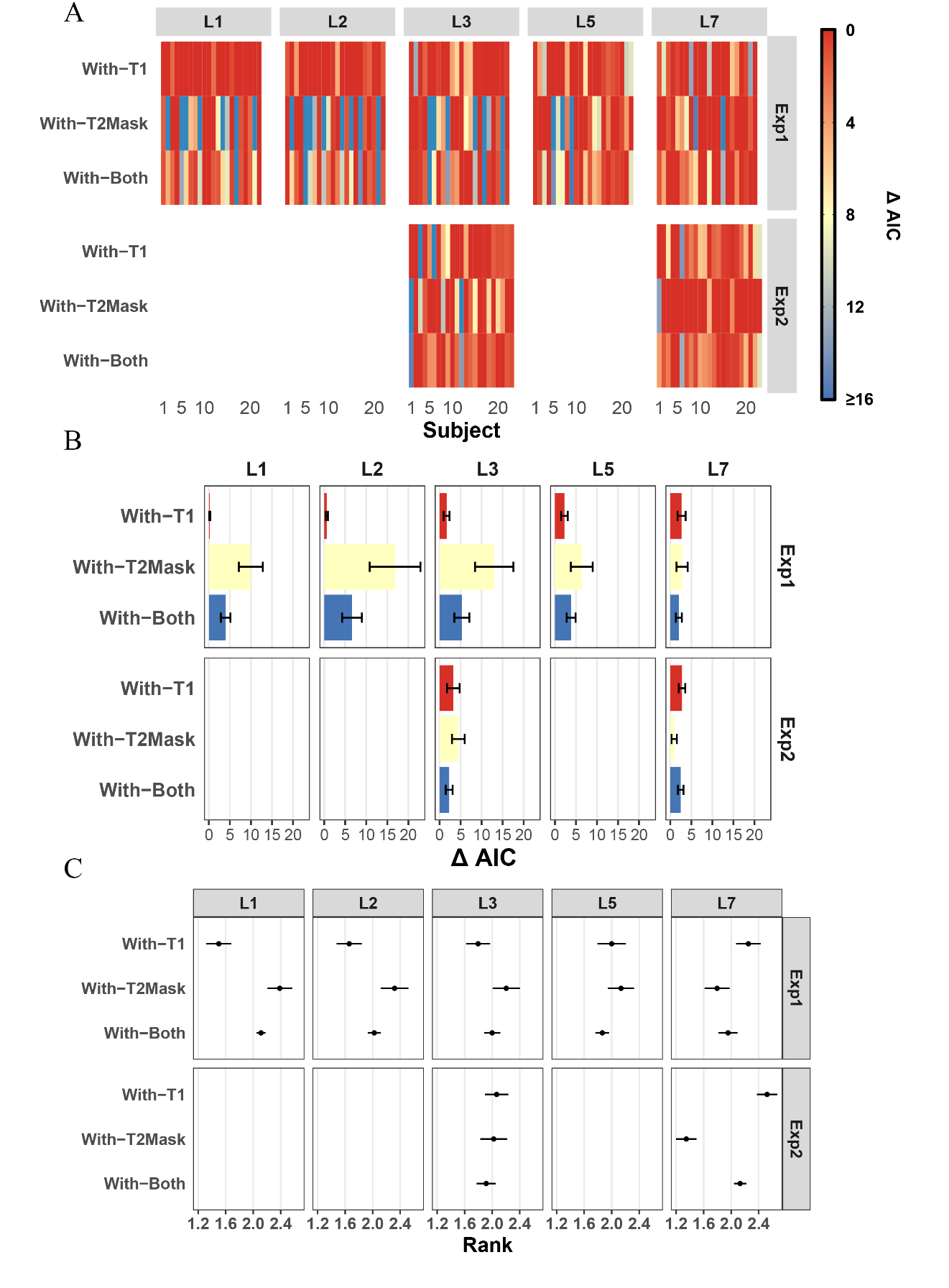
*Model comparison results for the Swap model and its variants.*

*Note.* (**A**) Model comparison results at the subject level, using the data from Tang et al. (2020) (**B**) Relative AIC values compared to that of the best-performing model, averaged across all subjects. (**C**) Average model ranking. Figure conventions follow Figure S1.
